# Imaging guided single-cell multiomics unveils shared autoreactive CD4+ T-cell responses in blood, locoregional lymph node and affected tissues of patients with systemic autoimmunity

**DOI:** 10.64898/2026.08.18.745406

**Authors:** Theodoros Ioannis Papadimitriou, Prashant Singh, Arjan van Caam, Xuehui He, Konnie M. Hebeda, Birgitte Walgreen, Elly Vitters, Pim Kloosterman, Laura J. A. Wingens, Klaas W. Mulder, Madelon Vonk, I. Jolanda M. de Vries, Peter van der Kraan, Ruben L. Smeets, Erik Aarntzen, Hans Koenen, Martijn A. Huynen, Rogier M. Thurlings

## Abstract

Systemic autoimmune connective tissue diseases (CTDs) are characterized by anti-nuclear antibodies, shared HLA-associated genetic risk, and frequent disease overlap, suggesting a central role for CD4+ T cells in pathogenesis. However, defining disease-driving CD4+ T-cell responses remains challenging due to their localization within lymphoid and affected tissues and the lack of approaches linking these responses to circulating counterparts. We combined [18F]-labeled thymidine PET/CT-guided tissue sampling, ex vivo antigen stimulation, and single-cell multiomics to characterize CD4+ T-cell responses in blood, PET-avid locoregional lymph nodes (LNs), and disease-affected tissues from patients with the immunologically distinct CTDs systemic sclerosis and Sjögren’s disease. PET-avid LNs from both diseases exhibited enhanced adaptive immune activity and contained an expanded population of interferon-stimulated gene (ISG)-expressing TRAIL+ CD4+ T cells. In Sjögren’s disease, active LNs and affected tissues harbored diverse effector CD4+ T-cell populations, including follicular and peripheral helper T cells and Th2/Th17 cells. In contrast, systemic sclerosis tissues lacked effector CD4+ T cells, while active LNs were enriched for naïve, regulatory, and TRAIL+ ISG CD4+ T cells. Antigen stimulation of peripheral blood mononuclear cells enriched for expanded effector CD4+ T-cell populations that shared activation profiles and clonal relationships with cells in LNs and affected tissues, many representing autoreactive antigen-specific T cells. TRAIL+ CD4+ T cells suppressed effector T-cell differentiation, autoreactive plasma cell generation, and autoantibody production in vitro, identifying a previously unrecognized immunoregulatory population. Together, this workflow enables comprehensive characterization of pathogenic and regulatory CD4+ T-cell responses across CTDs.

## Introduction

The systemic autoimmune connective tissue diseases (CTDs), systemic sclerosis (SSc), systemic lupus erythematosus, myositis and Sjögren’s disease (SjD), are debilitating diseases that frequently co-occur (1). They are characterized by a fluctuating disease course, systemic and local inflammatory responses, and organ- and connective tissue-specific symptoms (2). Although the etiology of these diseases remains poorly understood, loss of tolerance due to self-antigen recognition by autoreactive CD4+ T-cells is considered to play a central role (3). This is indicated by genetic studies confirming that polymorphisms in the HLA region confer the highest risk for development of CTDs. In addition, CTDs are characterized by circulating autoantibodies against nuclear antigens (ANA), and HLA class II alleles are associated with their development (4, 5).

In contrast, recent immunopathogenic and multi-omics studies that we and others performed indicate that the infiltration and activation of CD4+ T-cells in disease-affected tissues vary significantly across CTDs. SjD and SSc represent two CTDs at the opposite ends of this immunological spectrum. SjD is primarily characterized by chronic inflammation of salivary and lacrimal glands. These tissues exhibit a dense periductal infiltrate of effector CD4+ T-cell helper subsets (6, 7) and autoreactive B-cells (8). SSc on the other hand, is characterized by vasculopathy, autoimmune inflammation and fibrosis, leading to excessive extracellular matrix deposition in skin and internal organs such as heart, lungs and kidneys. In contrast with SjD, this involves a sparse perivascular immune cell infiltrate without an overt increase in CD4+ T-cells (9–11).

In contrast to experimental animal models of CTDs, the full spectrum of the aberrant adaptive immune response in human CTD —including priming, activation, and regulation of CD4+ T-cells in LNs, circulation, and affected tissues—remains unexplored. This is primarily due to the challenge of localizing disease-involved CD4+ T-cells, which are present in low numbers and reside in hard-to-access locoregional lymph nodes and disease-affected tissues. Additionally, there is a lack of imaging techniques capable of detecting and localizing LNs and affected tissues hosting an active autoreactive immune response. As a result, CD4+ T-cells have primarily been detected in blood circulation, limiting understanding of their generation in LNs and their role in disease-affected tissue pathology (12, 13). Advancements in imaging, such as positron emission tomography (PET), allow sensitive visualization and quantification of biomolecular processes simultaneously at the whole-body scale.

[^18^F]-labeled fluoro-2-deoxy-2-d-glucose ([^18^F] FDG) PET is routinely applied in clinical care to determine the activity of inflammatory diseases (14). However, glucose uptake is particularly increased in activated macrophages (15–17) and is less sensitive for detecting the activation of lymphocytes (29). [^18^F]-labeled 3⍰-fluoro-3⍰-deoxy-thymidine ([^18^F] FLT) detects thymidine incorporation during DNA synthesis phases of cell proliferation (18) and has been used to visualize active LNs in vaccination and oncology studies to guide sampling of LNs harboring an active adaptive immune response (19, 20).

Over the past decade high throughput detection of antigen-specific T-cells has been achieved by detection of T-cells expressing activation markers after ex vivo antigen-stimulation of peripheral blood mononuclear cells in the context of vaccination, infection (31), and more recently for autoimmune diseases (32–34). This approach, termed the activation-induced marker (AIM) assay, enables the sensitive and scalable identification of antigen-specific T cells without requiring prior knowledge of specific peptide–MHC restrictions. As such, it has become a widely adopted method for tracking antigen-specific T cell responses across diverse settings, allowing simultaneous interrogation of multiple epitopes and populations in a high-throughput manner without the need for MHC-restricted tetramers.

Here, we hypothesized that the differences in CD4+ T-cell infiltration and activation in disease-affected tissues of patients with SSc and SjD result from variations in the activation and regulation of CD4+ T-cell-mediated humoral immune responses originating in locoregional LNs. We used [^18^F] FLT PET imaging to identify lymph nodes containing proliferating lymphocytes. Followingly, we applied multimodal single-cell RNA sequencing combined with multiplex immunohistochemistry and ex vivo antigen-stimulation assays to define the properties of disease-involved CD4+ T-cell responses in blood, LNs and affected tissues.

## Results

### [^18^F] FLT PET scans in CTD patients distinguish lymph nodes harboring active adaptive immune response from quiescent lymph nodes

To understand if the extent of CD4+ T-cell infiltration and activation in CTD-affected tissues is related to the immune response in LNs, we undertook a multimodal single-cell approach to characterize CD4+ T-cell responses in the blood, affected tissues and locoregional LNs of patients with CTDs. We performed [^18^F] FLT PET scans to evaluate their usefulness in CTD patients for identifying LNs, harboring an active adaptive immune response in the proximity of disease-affected tissues. We selected treatment-naïve patients with SSc and SjD who had clinically active disease and were sero-positive for the ANA anti-Scl70 and anti-Ro60/La *(<u>Figure 1A, B</u>* and <u>*Supplemental Table 1*</u>). Anti-Scl70 and anti-Ro60/La ANA are associated with a severe disease course in SSc and SjD, respectively. LNs showing increased [^18^F] FLT PET uptake were considered as “positive” while those with no or low uptake were classified as “negative” *(<u>Figure 1C</u>)*. This imaging strategy guided the selective sampling of ultrasound-guided biopsies from positive and negative LNs. In patients with SjD, cervical LNs were biopsied *(<u>Figure 1D</u>)*, while in patients with SSc, LNs from disease-affected arms or legs were collected (*<u>Supplemental Figure 1</u>* and <u>*Supplemental Table 1*</u>). In addition, biopsies of disease-affected tissues were obtained from the same patients (salivary gland [SG] for SjD and skin for SSc).

**Figure 1.**
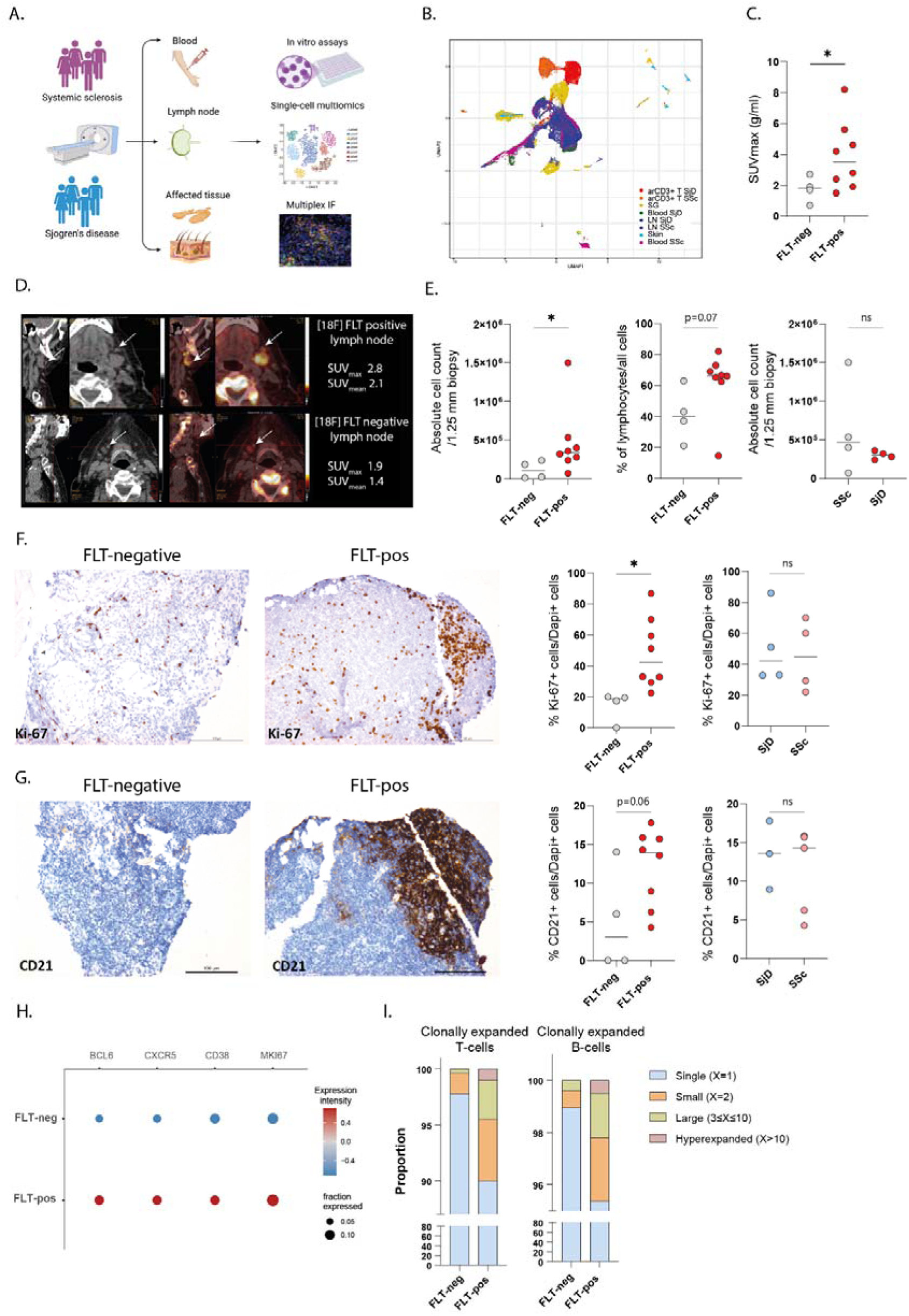
FLT PET scans allow identification of active lymph nodes containing clonally expanded T- and B-cells. **(A)** Overview of the experimental approach. Blood, lymph node (LN) and affected tissue (skin or salivary gland) samples from patients with SSc and SjD were isolated and processed for single-cell multimodal profiling (transcriptomics, proteomics, TCR/BCR sequencing). **(B)** UMAP displaying all cells recovered from SSc and SjD patients’ tissues, blood and ex vivo antigen-responsive (ar) CD3+ T-cells. Each subpopulation is illustrated with a different color. **(C)** SUV_max_ values between FLT-positive and FLT-negative LNs. **(D)** Representative [^18^F] FLT PET/CT scans from the cervical area of a patient with SjD (n=4) exhibit with arrows detection, among the salivary gland (SG) draining lymph nodes, of a FLT-positive and a FLT-negative LN based on high or low [^18^F] FLT signal, represented with SUV_mean_ and SUV_max_ values (g/ml). **(E)** Absolute cell counts and % of CD45+ lymphocytes among all cells of each biopsy comparing FLT-positive versus FLT-negative LNs and FLT-positive LNs of patients with SjD versus SSc. Representative images of KI-67 **(F)** and CD21 **(G)** immunohistochemistry staining of FLT-negative and FLT-positive LN of a patient with SjD, accompanied by quantification of KI-67+ or CD21+ cells (n=4 FLT-negative, 4 FLT-positive SSc, 4 FLT-positive SjD LNs). **(H)** 2D dot plots comparing the gene expression of the depicted genes between FLT-positive and FLT-negative LNs. (I) Comparison of the proportion of clonally expanded T-cells and B-cells between FLT-negative/positive LNs. Each symbol and each horizontal line in (C), (E), (F), (G) represents one donor and mean value respectively. Statistical differences: two-tailed unpaired t test with significance set at *p<0.05.

Immunohistochemical analysis confirmed earlier findings that salivary glands affected by SjD contained a large CD4+ T-cell infiltrate, whereas skin affected by SSc exhibited only a sparse CD4+ T-cell infiltrate (Supplemental Figure 2A). To confirm that FLT-PET guidance successfully identified LNs with proliferating lymphocytes, we analyzed lymphocyte retrieval from cell suspensions and examined immunohistochemical markers in tissue biopsies. In FLT-positive LNs (n=4 SSc and n=4 SjD), a higher number of total cells per biopsy and a higher fraction of lymphocytes were observed compared to FLT-negative LNs (n=2 SSc and n=2 SjD; fewer due to sampling constraints related to anatomical localization and bleeding risks) (median yield 340,000 cells versus 105,162 cells respectively, with 66% versus 40% lymphocytes) *(<u>Figure 1E</u>)*. There was no difference in the total number of cells in FLT-positive LNs between SSc and SjD patients *(<u>Figure 1E</u>)*. The number of proliferating cells, as assessed by Ki-67 immunohistochemistry, was also higher in FLT-positive compared to FLT-negative LNs (48% of Ki-67+ cells compared to 14%) and did not differ between SSc and SjD patients *(<u>Figure 1F</u>)*. To determine whether cell proliferation was driven by APC-mediated T-cell and B-cell activation, we analyzed the extent of the adaptive immune response, by assessing CD21 expression, a marker for mature B-cells and dendritic cells (DCs) and by T-cell and B-cell clonality analysis. FLT-positive LNs contained an elevated number of CD21+ mature B-cells/DCs in comparison to FLT-negative LNs *(<u>Figure 1G</u>)*. Additionally, gene expression analysis revealed increased levels of *MKI67* and the follicular markers *CXCR5, BCL6* and *CD38* in FLT-positive LNs *(<u>Figure 1H</u>).* Finally, clonality analysis demonstrated that FLT-positive LNs contained more expanded T- and B-cell clones compared to FLT-negative LNs *(<u>Figure 1I</u>)*. In summary, these findings indicate a more active adaptive immune response in FLT-positive LNs in both SSc and SjD (hereafter termed as active LNs).

### Active lymph nodes in SjD contain lymphocyte clusters, while in SSc these contain dispersed lymphocytes

To understand the differences in activation and regulation of adaptive immune responses within active LNs of SSc and SjD patients, we analyzed the type and interaction of adaptive immune cell subsets in the biopsies from active LNs, SjD affected SGs and SSc affected skin. We first undertook a qualitative approach, using a semi-quantitative score to assess the level of T/B-cell clustering within APC rich lymphocyte areas, identified by the markers CD3 (T-cells), CD79A/CD20 (B-cells) and CD1c/CD21 (APCs/mature B-cells). In SjD affected SGs, we observed large and germinal center (GC)- like organized aggregates of APCs, T-cells and B-cells. Similar to SjD affected SGs, active LNs of SjD patients contained large and GC-like organized aggregates of APCs, T-cells and B-cells. In contrast, SSc affected skin contained small aggregates of APCs and T-cells, and no B-cells (Supplemental Figure 2B). In SSc active LNs, T-cells, B-cells and APCs were dispersed throughout the biopsy without forming evident GC-like structures *(<u>Figure 2A, B</u>).* These findings were further confirmed by multiplex immunofluorescence staining *(<u>Figure 2C, D</u>)*.

**Figure 2.**
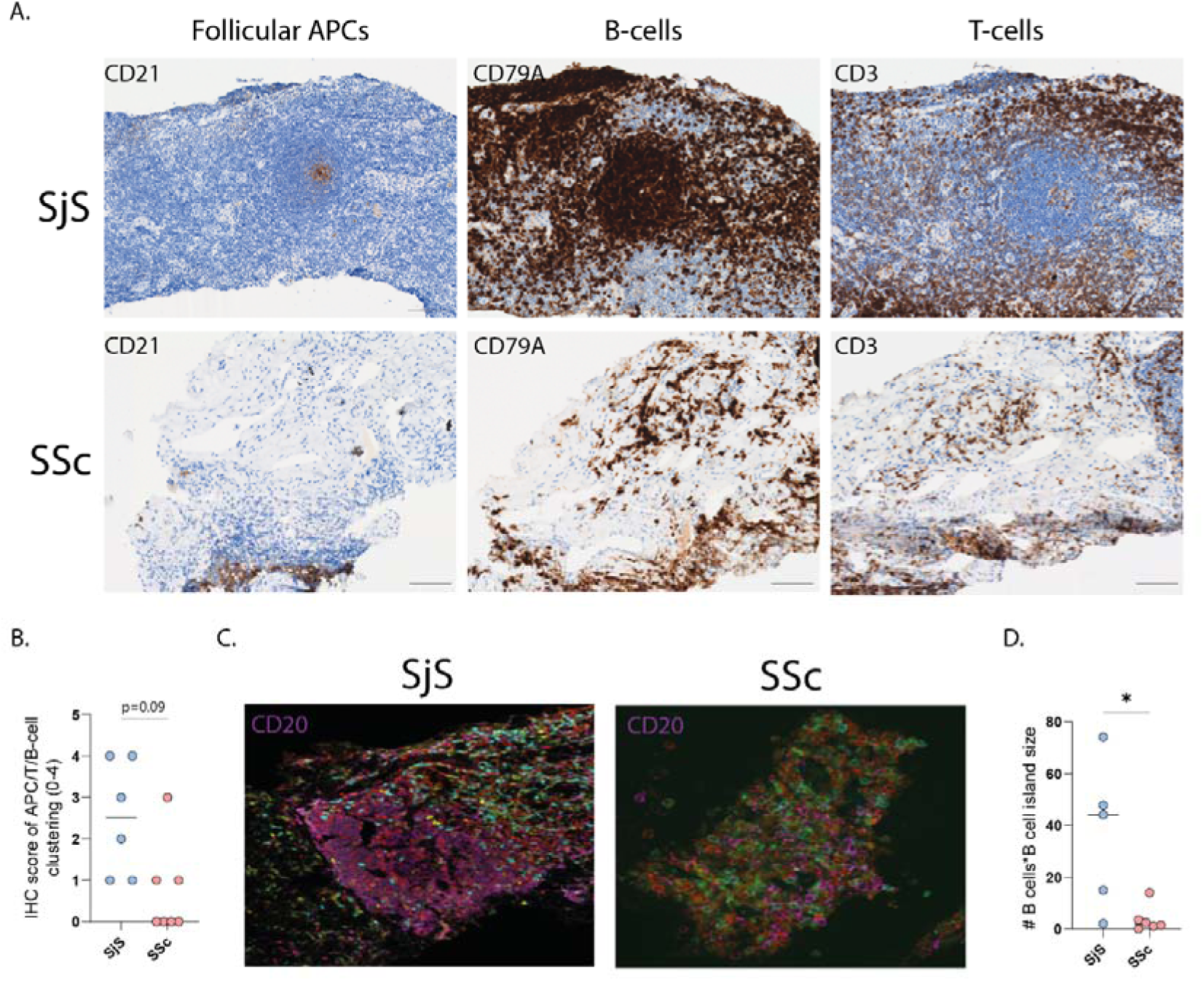
APC-T-B-cell interaction in SSc lymph nodes is limited compared to more organized and robust in SjD. **(A)** IHC stainings of representative SjD (n=6) and SSc (n=7) active LNs for CD21, CD79A and CD3. Scale bars are 100 μm. **(B)** Semi-quantitative IHC score (values range from 0-4) for the quantification of the level of APC-T-B-cell clustering (n=6 SjD, n=7 SSc). **(C)** Representative Immunofluorescent images and **(D)** quantification of the number of B-cell islands multiplied by total B-cells per biopsy in SjD (n=5) and SSc (n=6) patients’ active LNs. The following markers were used for the immunofluorescent staining; CD20 (purple), CD3 (red), CD56 (yellow), FOXP3 (green), CD8 (cyan). Each symbol and each horizontal line in (B), (D) represents one donor and mean value respectively. Statistical differences: two-tailed unpaired t test with significance set at *p<0.05.

### Active lymph nodes of SjD contain a pleiotropic CD4+ T-cell effector response, while in SSc these contain regulatory and CD4+ TRAIL+ ISG T-cells

Subsequently we focused on the activation and function of the CD4+ T-cells in active LNs using single-cell multi-omic analysis of the T-cells of the same LNs as above. We performed unsupervised whole transcriptome clustering of isolated T-cells, focusing on CD4+ T-cell populations *(<u>Figure 3A</u>)*. CD4+ T-cell clusters were identified based on differentially expressed genes, differentially expressed proteins, expression of canonical subset markers, and joint density of multiple features *(<u>Figure 3B, C</u>* and *Supplemental Figure 3A, B)*. This analysis identified eight distinct CD4+ T-cell clusters: naïve, central memory (CM), regulatory (Tregs), TRAIL+ interferon-stimulated genes (TRAIL+ ISG), T-helper (a mix of Th2 and Th17 cells), activated (CD4 activ), T-cell helper providing B-cell help (a mix of extrafollicular helper and follicular helper T-cells [Tfh/ef]) and a small cluster of unknown (CD4 uk) identity. Compared to non-active LNs, SjD active LNs contained a significantly lower amount of naïve CD4+ T-cells and a higher percentage of effector CD4+ T-cells, including increased T-helper, T-fh/ef, CD4+ activ and TRAIL+ CD4+ ISG T-cells *(<u>Figure 3D</u>)*. In contrast, SSc active LNs did not significantly differ in the fraction of naive T-cells compared to the quiescent ones, and CD4+ T-cells mainly consisted of regulatory T-cells and TRAIL+ CD4+ ISG T-cells *(<u>Figure 3D</u>).* Finally, we analyzed the expanded TCR clones in each CD4+ T-cell cluster. A small proportion of ∼7% of CD4+ T-cells was expanded in active LNs. T-helper cells and Tfh/ef exhibited the highest proportion of expanded T-cell clones, and their clonal expansion was more pronounced in SjD than in SSc *(<u>Figure 3E</u>).* Collectively, these results indicate that active LNs in both diseases show signs of an active memory CD4+ T-cell response. Only SjD active LNs harbor a B-cell helper T-cell response, consistent with the observed histological APC/B/T-cell interaction clusters in SjD. In contrast, SSc active LNs are characterized by a TRAIL+ CD4+ ISG T-cell population, without an increased Treg/Teff ratio (*<u>Supplemental Figure 3C</u>*).

**Figure 3.**
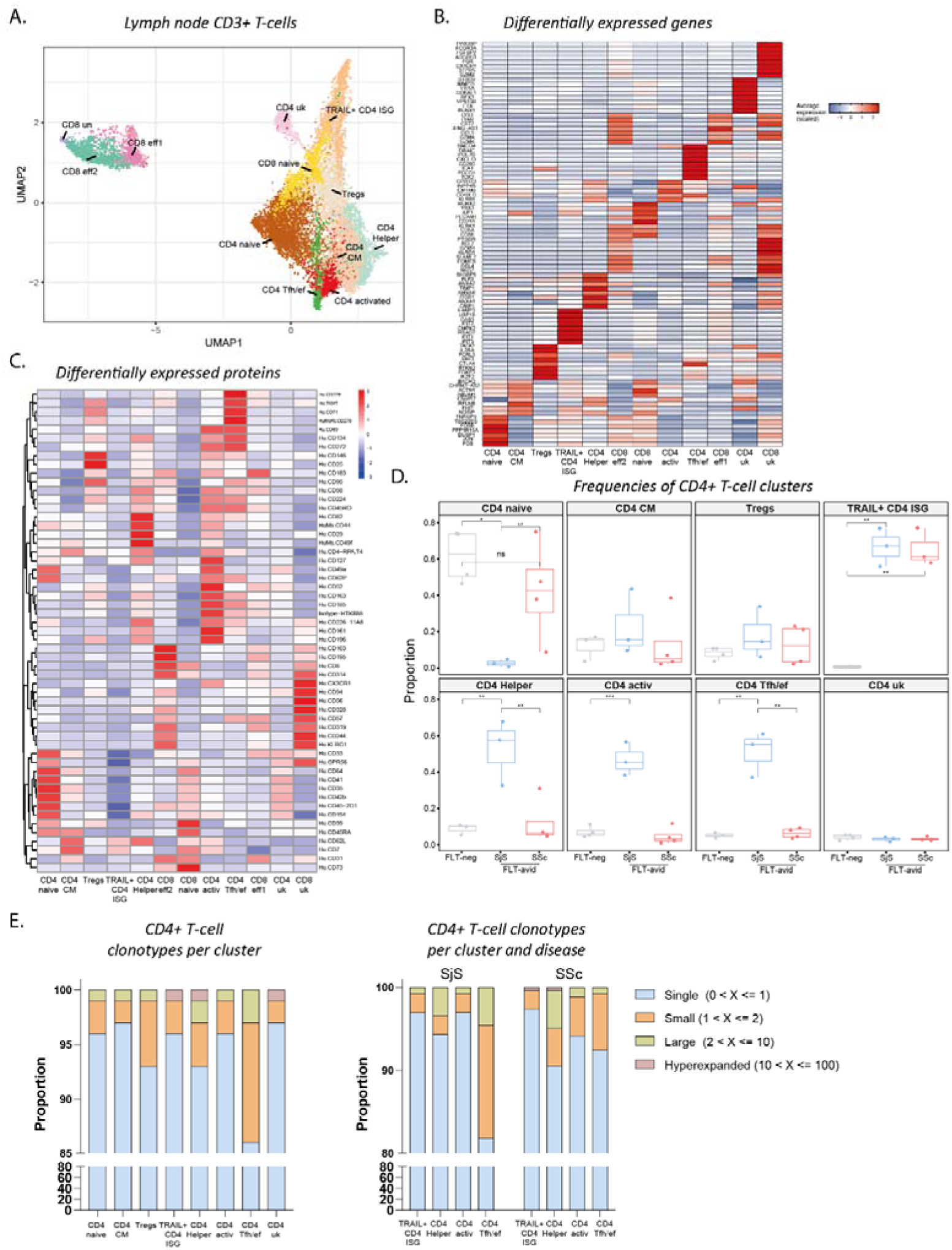
Single-cell analysis of active LNs reveals a more evident B helper CD4+ T-cell response in SjD compared to SSc. **(A)** UMAP plot of CD3+ T-cell subsets from patients’ FLT negative and positive LNs. **(B)** Heatmap illustrating the top differentially expressed genes in each distinguished T-cell cluster. **(C)** Heatmap exhibiting the top differentially expressed proteins for each transcriptionally distinct T-cell cluster. **(D)** Frequencies of CD4+ T-cell clusters in the FLT negative (n=4) compared to FLT positive LNs of patients with SjD (n=4) and SSc (n=4). Values are presented as the percentage variation in cell counts. Statistical analysis was conducted using the Wilcoxon test, with correction for multiple comparisons, *q<0.05, **q<0.01, ***q<0.001. **(E)** Distribution (proportion) of clonally expanded T-cells based on clonotype size for each CD4+ T-cell cluster. For the clusters of TRAIL+CD4 ISG, CD4 Helper, CD4 activ and Tfh/ef clonal distribution between SjD and SSc is also illustrated to the right.

### A distinct T helper subtype of CD4+ TRAIL+ ISG T-cells is uniquely present in the extrafollicular areas of patients’ active lymph nodes

To gain more insight into the identity of TRAIL+ CD4+ ISG T-cells, we analyzed their transcriptome, proteome and tissue localization using single-cell RNA sequencing, CITE-seq and multiplex immunofluorescence. TRAIL+ CD4+ T-cells have been studied for their dual roles in inducing cell death and regulating immune responses via binding of TRAIL to its receptors DR4 and DR5. CD4+ T-cells with ISG expression have been identified in single-cell analyses across various conditions, including viral infections, lupus, and rheumatoid arthritis (21–25). Here, the detected CD4+TRAIL+ISG T-cells were characterized by the expression of the death receptor TRAIL (*TNFSF10*) and a prominent type I ISG signature, including *MX1, IFIH1, OASL,* and *IFITs (<u>Figure 4A</u>, <u>Supplemental Figure 3B</u>)*. Additionally, CD4+TRAIL+ ISG T-cells expressed type 1 interferon receptors (*IFNAR1/2*) and intracellular double stranded RNA receptors (*DDX58, DHX58* and *IFIH1*) but did not express genes that mediate detection of intracellular DNA (cGAS or STING) *(Supplemental Figure 3B).*

**Figure 4.**
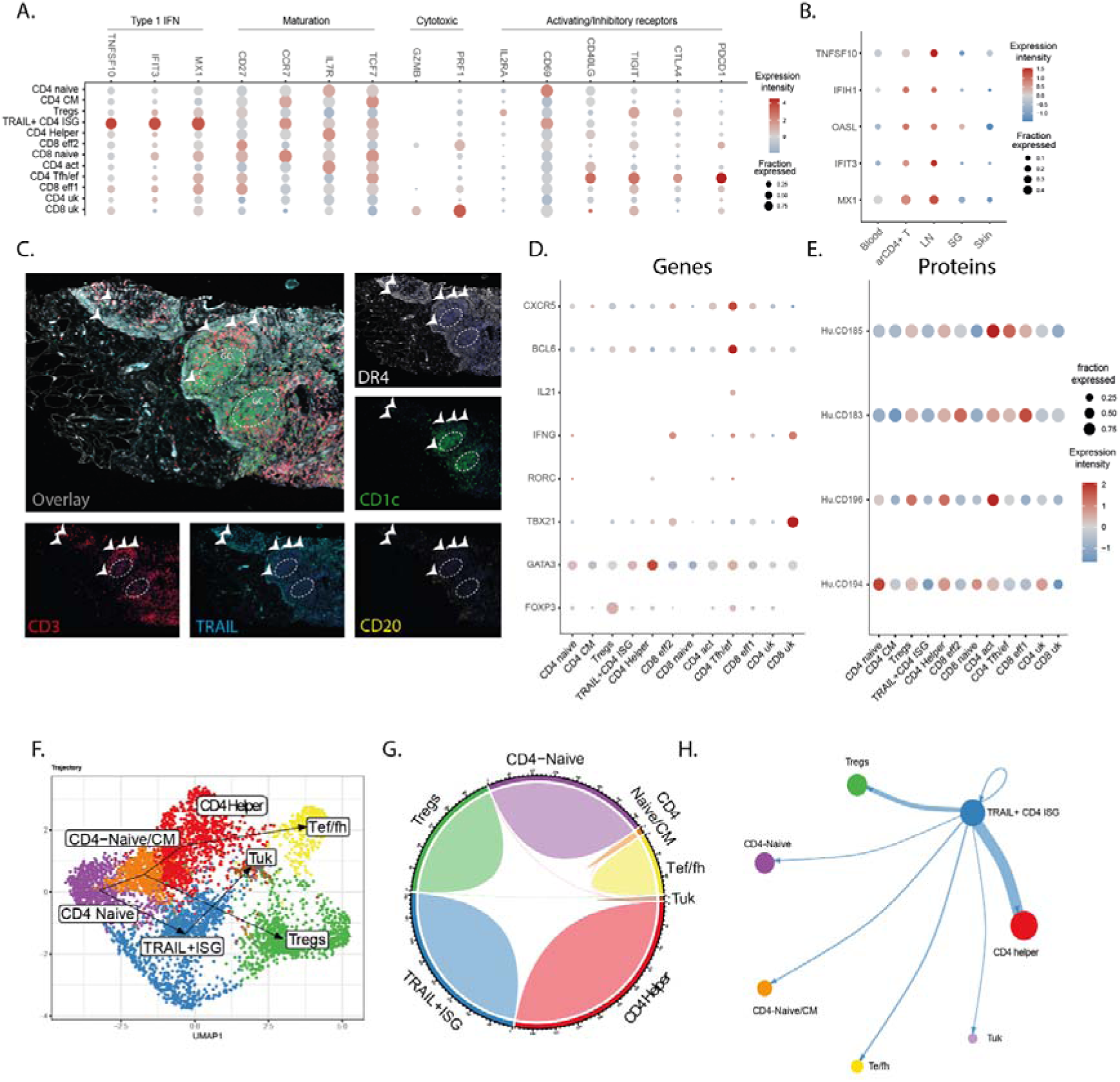
CD4+ TRAIL+ ISG T-cells render a distinct naive-like T helper subtype that is uniquely present in extrafollicular sites of patients’ reactive LNs. 2D dot plots comparing gene expression of selected genes **(A)** per cluster **(B)** between blood, active LNs and arCD4+ T-cells. **(C)** Multicolor immunofluorescence staining of the depicted markers to visualize the spatial mapping of CD4+ TRAIL+ T-cells within reactive LNs. Scale bars are 100 μm. 2D dot plots comparing **(D)** gene and (E) protein expression of selected CD4+ T helper markers between CD4+ T-cell clusters. (F) Single-cell developmental trajectories of LN CD4+ T-cell clusters. **(G)** Cord diagram showing the TCR clonal overlap between the LN CD4+ T-cell clusters. **(H)** Graphical representation of cell-cell edges (interactions) of TRAIL+ ISG CD4+ T cells with the rest CD4+ T-cell clusters, exhibiting high likelihood and specificity of interaction of TRAIL+ ISG CD4+ T cells with CD4+ effector T-cells (thick blue line).

Regarding their maturation stage, these cells exhibited a naïve-like phenotype (high *SELL, CCR7*; moderate *CD27, IL7R*) without markers of cytotoxicity, T-cell activation/exhaustion or follicular localization (low *CXCR5, BCL6*) *(<u>Figure 4A</u>, <u>Supplemental Figure 3B</u>)*. This indicates a relatively quiescent extrafollicular state. Comparative analysis between tissues confirmed that these cells were not observed in affected tissues, but selectively found in patients’ active LNs *(<u>Figure 4B</u>, <u>Supplemental Figure 4</u>).* Immunohistochemical localization studies showed that they occupied extrafollicular zones in active LNs, adjacent to germinal center-like structures in SjD or scattered near DR4+ APCs in SSc *(<u>Figure 4C</u>, <u>Supplemental Figure 4</u>)*. The TRAIL+ CD4+ ISG T-cell cluster in active LNs was transcriptionally and phenotypically distinct from Th1, Th2, Th17, Tfh, and Treg subsets, lacking apparent gene expression of *TBX21, GATA3, RORC, BCL6* and *FOXP3 (<u>Figure 4D</u>)* and protein expression of CXCR3 (CD183), CXCR5 (CD185), CCR4 (CD194) and CCR6 (CD196) compared to the CD4 effector and Treg clusters *(<u>Figure 4E</u>).*

Pseudo-time analysis of CD4+ T-cell differentiation positioned TRAIL+ CD4+ ISG T-cells along a distinct trajectory, separate from helper and regulatory T-cells, originating from naïve CD4+ T-cells *(<u>Figure 4F</u>*, *<u>Supplemental Figure 5A-C</u>*). Of note, TCR clonal overlap analysis showed minimal overlap with helper subsets and no overlap with Tregs, further supporting a unique identity *(<u>Figure 4G</u>)*. Differential dynamic gene expression analysis highlighted type I IFN-stimulation responsive genes (*STAT1, MX1, ISG15*), T-cell activation markers (*CD69*), and TCR signaling genes (*LY6E*) as key contributors to the transition of CD4+ naïve T-cells to the TRAIL+ ISG phenotype (Supplemental Figure 5D). Cell-cell communication analysis using CellChat, a computational framework that infers intercellular signaling networks based on known ligand–receptor interactions, showed that TRAIL+ CD4+ T-cells exhibited a robust interaction with CD4+ effector T cells *(<u>Figure 4H</u>)*.

To gain insight in differences in TRAIL+ ISG CD4+ T-cell activation between active CTD LNs and physiological settings we merged our scRNAseq T-cell dataset with a public dataset of T-cells from tonsils. Notably, both the frequency of TRAIL⁺ ISG CD4⁺ T cells and the intensity of their ISG signature were elevated in LNs compared to tonsils (22% vs 1.2%) *(Supplemental Figure 6A-D)*. TRAIL+ ISG CD4+ T-cells were significantly clonally expanded in SSc reactive LNs, compared to SjD reactive LNs and tonsils *(Supplemental Figure 6E)*. Furthermore, LN TRAIL+ CD4+ ISG T-cells expressed genes linked to immunoregulatory pathways *(Supplemental Figure 6F)*, whereas in tonsils, their gene profile was more viral infection-related. In summary, CD4+TRAIL+ ISG T-cells are a uniquely expanded CD4+ T-cell subset in active LNs of CTD patients, especially SSc, with a distinct extrafollicular lymph node localization and mixed gene expression markers of quiescence and activation.

### Expanded CD4+ T cell clones from active LNs and disease-affected tissues recirculate through blood, where they can be detected as antigen-stimulus responsive CD4+ T-cells

Because of the challenge to obtain disease-involved T-cells from active LNs and disease-affected tissues, it would be beneficial to identify circulating counterparts of tissue-resident T-cells. To confirm our findings in a larger patient cohort and to obtain insight if expanded CD4+ T-cell clones in active LNs and disease-affected tissues recirculate in blood, we analyzed activated CD4+ T-cells (arCD4+ T-cells) of SjD and SSc patients using an activation induced marker (AIM) assay. For this, we pulsed PBMCs of SSc (n=45), SjD (n=38) patients and healthy controls (n=30) with protein autoantigens (Ro60/La for SjD, Scl70 for SSc) and detected antigen stimulation-responsive CD4+ T-cells based on their upregulation of CD25 and OX40 (CD134) *(<u>Figure 5A, B</u>)*. We observed a robust fraction of CD4+ T-cells responsive to autoantigen stimulation in SjD, compared to a limited fraction in SSc blood (*<u>Figure 5C</u>*). Ro60/La arCD4+ T-cells were detected in higher numbers in healthy controls compared to Scl70-arCD4+ T-cells, but these were predominantly naive compared to predominantly memory in SjD patients (*<u>Figure 5D</u>*).

**Figure 5.**
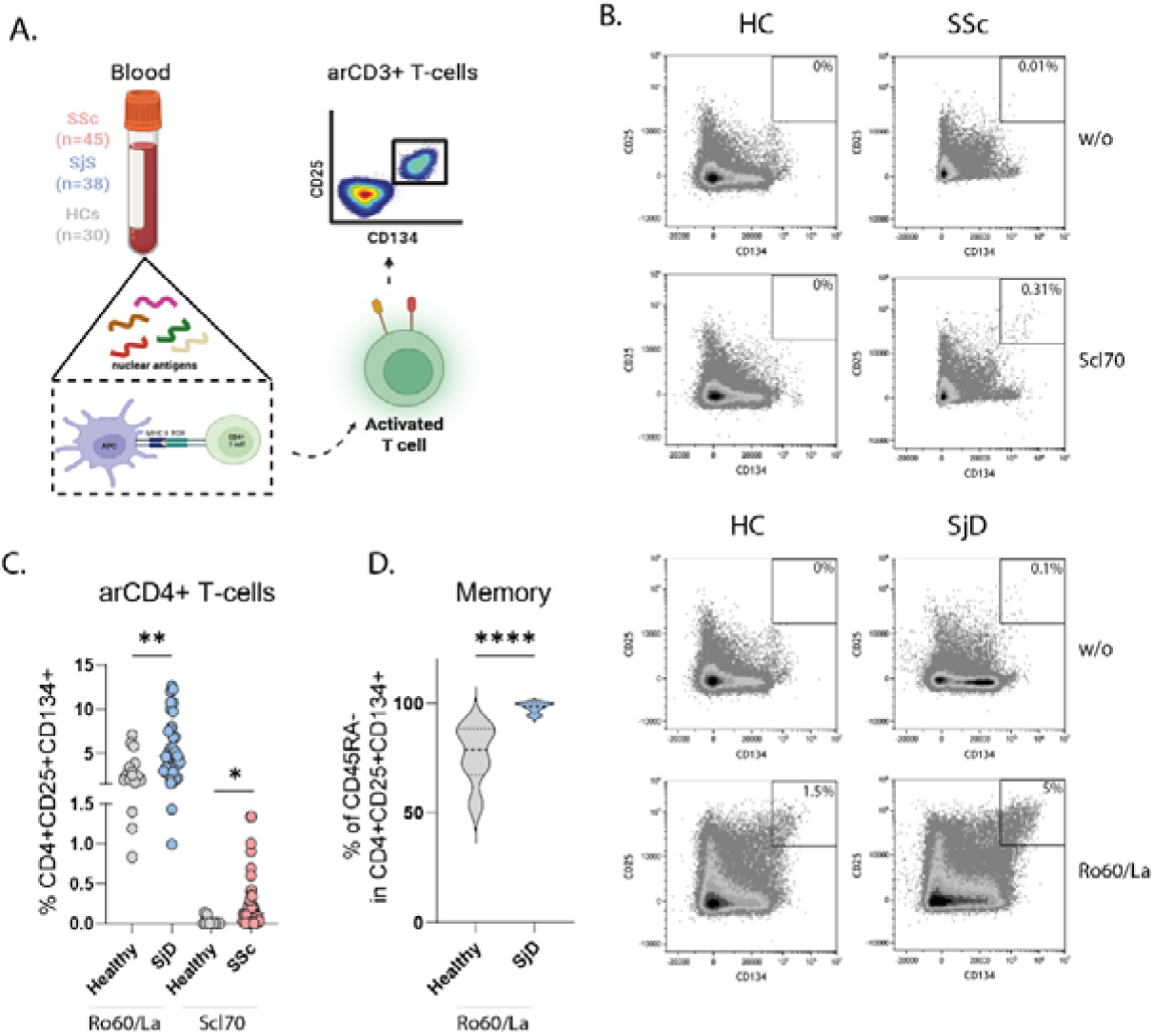
Detection and characterization of antigen-responsive CD4+ T-cells with the AIM assay. **(A)** Experimental workflow for the detection and characterization of arCD4+ T-cells in patients with SjD and SSc and healthy donors (HC). **(B)** Representative flow cytometry plots exhibiting the detection of arCD3+ T-cells identified as CD4+CD25+CD134+ after 16 hours of incubation of PBMCs with or without anti-nuclear antigen (w/o, [Scl70 in 45 SSc patients, Ro60/La in 37 SjD]). **(C)** Frequencies of arCD3+ T-cells in HCs (n=30), and patients with SjD (n=37) and SSc (n=45). **(D)** Percentage of memory (CD45RA-) CD4+ T-cells within CD4+CD25+CD134+ T-cells in HCs (n=6) and SjD patients (n=17). Each symbol and each horizontal line in (C) represent one donor and mean value respectively. Statistical differences: two-tailed unpaired t test with significance set at *p<0.05, **p<0.01, ****p<0.0001.

Interestingly, in 2 HLA-matched SjD patients 60% of Ro60-stimulated arCD4+ T-cells overlapped with Ro60-MHC class II tetramer sorted T-cells (Supplemental Figure 7A-C). Of note, combining the AIM assay with Ro60-MHC class II tetramer staining, showed that AIM+ arT-cells contained 11-times more Ro60+ T-cells compared to their AIM-counterpart. Furthermore, their induction could be abrogated by MHC class II blockade, suggesting an antigen-specific nature of arCD4+ T-cells (Supplemental Figure 8). This notion is further supported by in vitro experiments in which arCD4⁺ T cells were single-cell sorted, expanded, and subsequently re-stimulated with autologous APCs pulsed with the respective antigen, demonstrating that arCD4⁺ T cells responded to restimulation by producing IFNγ.

To investigate the link between arCD4+ T-cells in blood and CD4+ T-cells in active LNs and disease-affected tissues, we performed single-cell RNA and TCR sequencing of sorted arCD4+ T-cells along with CD4+ T-cells from blood, paired affected tissues and LNs from the same patients (n=5 SSc and n=4 SjD *(<u>Figure 6A</u>)*. ArCD4+ T-cell clones exhibited higher clonal expansion compared to total CD4+ T-cells from blood *(<u>Figure 6F</u>)*. Almost all arCD4+ T-cells were memory cells (SjD; 99%, SSc; 99%) compared to ∼48% memory T-cells in peripheral blood (SjD; 44%, SSc; 51%) (*<u>Figure 6G-H</u>*). Similarly, all clonal counterparts of arCD4+ T-cell clones in active LNs/affected tissues were memory T-cells, compared to 41% of all T-cells in active LN being memory T-cells (SjD; 41%, SSc; 41%) and 82% in affected tissues (SjD SG; 72%, SSc skin; 91%). Eighteen percent (2,300/13,000 cells) of arCD4+ T-cells were part of clones that were also detected in tissue biopsies. The number of arCD4+ T-cell clones that were also detected in LN (SjD; 104, SSc; 38) and affected tissue biopsies (SjD SG; 42, SSc skin; 3) was larger compared to the number of shared clones between T-cells in blood and LNs (SjD; 14, SSc; 8), and between T-cells in blood and affected tissues (SjD SG; 10, SSc skin; 1) *(<u>Figure 6B</u>)*, Strikingly, while only 7% of CD4 T-cells in active LNs were clonally expanded, 35% of tissue residential clonal counterparts of arCD4 T-cells were expanded *(<u>Figure 3E</u>, <u>6E</u>)*. Together, this indicates that this assay enriched for circulating counterparts of expanded memory T-cell clones from active LNs and affected tissues, including a considerable proportion of autoreactive T-cells, allowing for phenotypic characterization and functional studies of disease-involved T-cell subsets.

**Figure 6.**
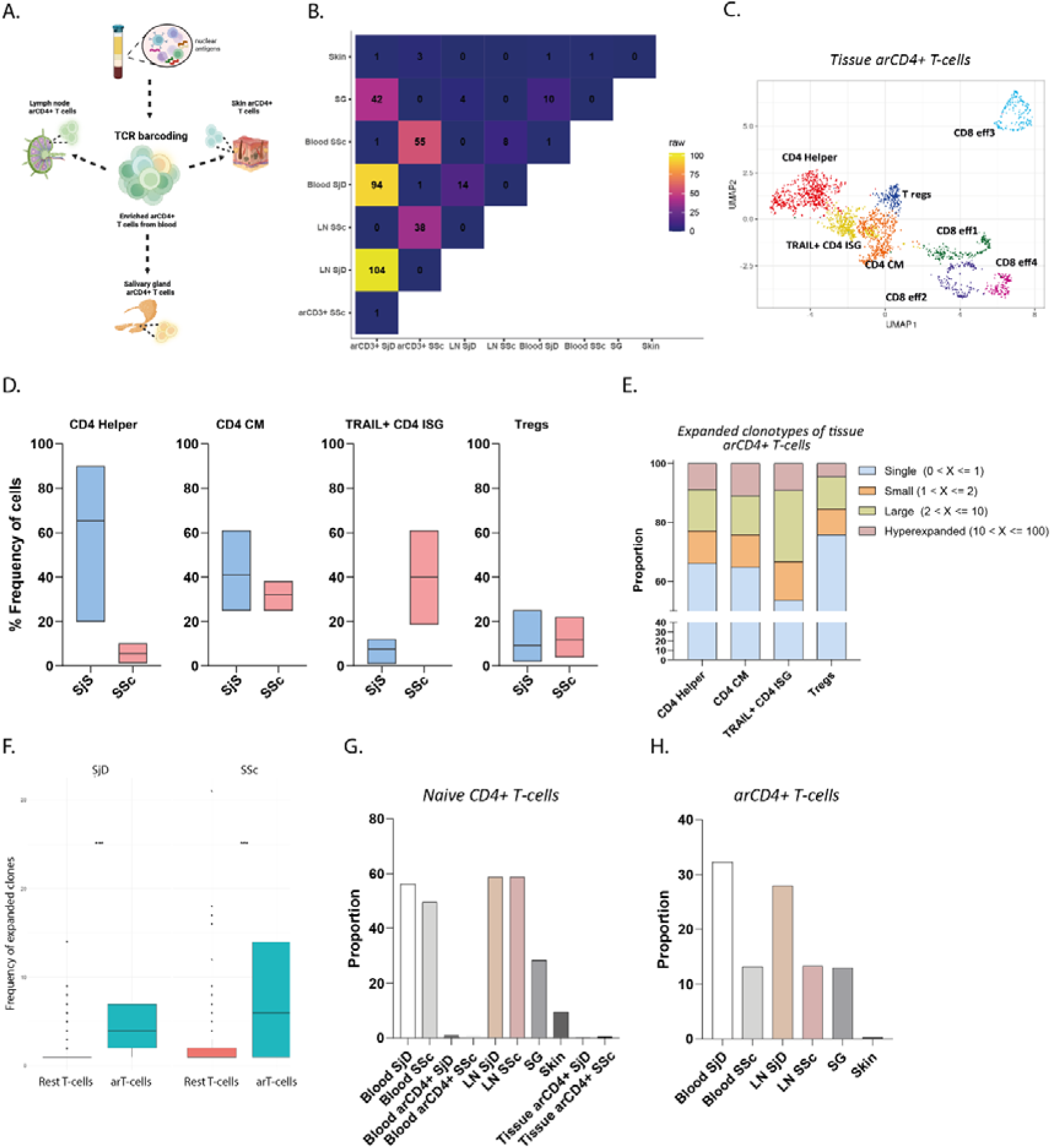
arCD4+ T-cells in SjD active lymph nodes and SGs exhibit robust effector functions, whereas in SSc, they display regulatory phenotypes. **(A)** Experimental workflow exhibiting the workflow used to detect and characterize arCD4+ T-cells in patients’ active LNs (n=4 SSc, n=4 SjD), skin (n=2) and salivary glands (n=3). **(B)** Morisita overlap quantification of the number of matching T-cell lineages among different tissues and between blood enriched arCD4+ T-cells and T-cells in tissues. **(C)** UMAP of arCD3+ T-cells in patients’ active LNs and affected tissues accompanied by **(D)** Cell frequency of arCD4+ subpopulations between SjD and SSc. **(E)** Proportion of clonal lineages by clonal size within each cluster of arCD4+ T-cells. **(F)** Frequency of expanded T cell clones between ar T cells and the rest (non-ar) T cells for both SjD and SSc. Tissue distribution of **(G)** naïve CD4+ T-cells and (H) arCD4+ T-cells.

Antigen-stimulus responsive CD4+ T-cell clones shared between blood, active lymph nodes and affected tissues display pleiotropic effector functions in SjD, whereas in SSc they display a regulatory and TRAIL+ ISG phenotype, coinciding with fewer arCD4+ T-cell clones that migrate in blood and SSc-affected skin.

To gain more insight in tissue-residential arCD4+ T-cells and circulating counterparts, we examined the presence and phenotype of arCD4+ T-cell clones in disease affected tissues, active LNs and blood. ArCD4+ T-cell clones were highly present in SjD SG, but none were found in SSc affected skin. Similarly, arCD4+ T-cell clones were more abundant in SjD blood and active LNs, compared to SSc blood and active LNs *(<u>Figure 6E</u>)*. In SjD, the affected SGs, active LNs and blood contained a large fraction of effector arCD4+ T-cell clones, that displayed a diverse CD4 T-helper phenotype that included Tfh, Tef, Th2, and Th17 cells *(<u>Figure 6F, G</u>)*. In contrast, arCD4+ T-cell clones in SSc LNs and blood predominantly exhibited a central memory (CM), TRAIL+ ISG, and Treg phenotype *(<u>Figure 6G</u>)*. The Tregs in SSc partly expressed markers of follicular Tregs (*FOXP3* and *BCL6*) and partly of extrafollicular Tregs (*FOXP3* and *PRDM1*). Circulating arCD4+ T-cells were clonal counterparts of the most expanded CD4+ T-cell clones in tissues for all T-cell subsets in both diseases *(<u>Figure 3E</u>, <u>6G</u>)*. Strikingly, the TRAIL+CD4+ISG T-cell subset exhibited the largest clonal expansion compared to other arCD4+ T-cell clones *(<u>Figure 6G</u>)*. While only 3% of TRAIL+CD4+ISG T-cells in LN were clonally expanded *(<u>Figure 3E</u>)*, 48% of the TRAIL+CD4+ISG clonal counterparts of circulating TRAIL+ arCD4+ ISG T-cells were expanded. Finally, the ratio of arCD4+ Tregs versus effector arCD4+ T-cells was higher in SSc compared to SjD *(<u>Figure 6D</u>)*. Collectively, these findings show that in SjD patients, arCD4+ T-cell clones are shared between blood, LNs and affected tissues, and entail a pleiotropic effector responses. In contrast, SSc is characterized by arCD4+ T-cell clones expressing a regulatory, and TRAIL+ ISG phenotype, coinciding with the generation of a limited arCD4+ T-cell effector response and fewer arCD4+ T-cell clones that migrate in blood and disease-affected tissues.

### SjD blood features more antigen-stimulus responsive CD4+ T-cells, displaying signs of B-cell help, compared to SSc where antigen-stimulus responsive CD4+ T-cells express TRAIL

To validate our findings and to render convenient read-outs for functional experiments, we analyzed if the single-cell multi-omic findings on arCD4+ T-cells in blood could be replicated and validated using flow cytometry in a larger patient cohort (45 SSc, 38 SjD and 30 HCs). With flow cytometry the arCD4+ T-cell response in blood showed higher abundance and an elevated CD4+/CD8+ T-cell ratio in SjD compared to SSc patients *(<u>Figure 5A</u>, <u>7A</u>)*. Also, arCD4+ T-cells in SjD exhibited elevated expression of the B-cell help markers CD40L, IL21 and were enriched for a CXCR5+PD-1+ICOS+ Tfh phenotype *(<u>Figure 7B, C</u>)*. In contrast, arCD4+ T-cells in SSc blood showed elevated TRAIL expression, while the expression of the death receptor ligand FASL did not differ between the diseases *(<u>Figure 7D</u>)*. TRAIL expression was restricted to memory arCD4+ T-cells *(<u>Figure 4A</u>)*. It was absent in unstimulated PBMCs, and in arCD4+ T-cells after MHC class II blocking *(<u>Figure 7E</u>)*. Taken together, flow cytometric analysis confirms that circulating arCD4+ T-cells are more prevalent in SjD compared to SSc and that these show a similar polarization as expanded CD4+ T-cell clonal counterparts of arCD4+ T-cells in active LNs of these conditions.

**Figure 7.**
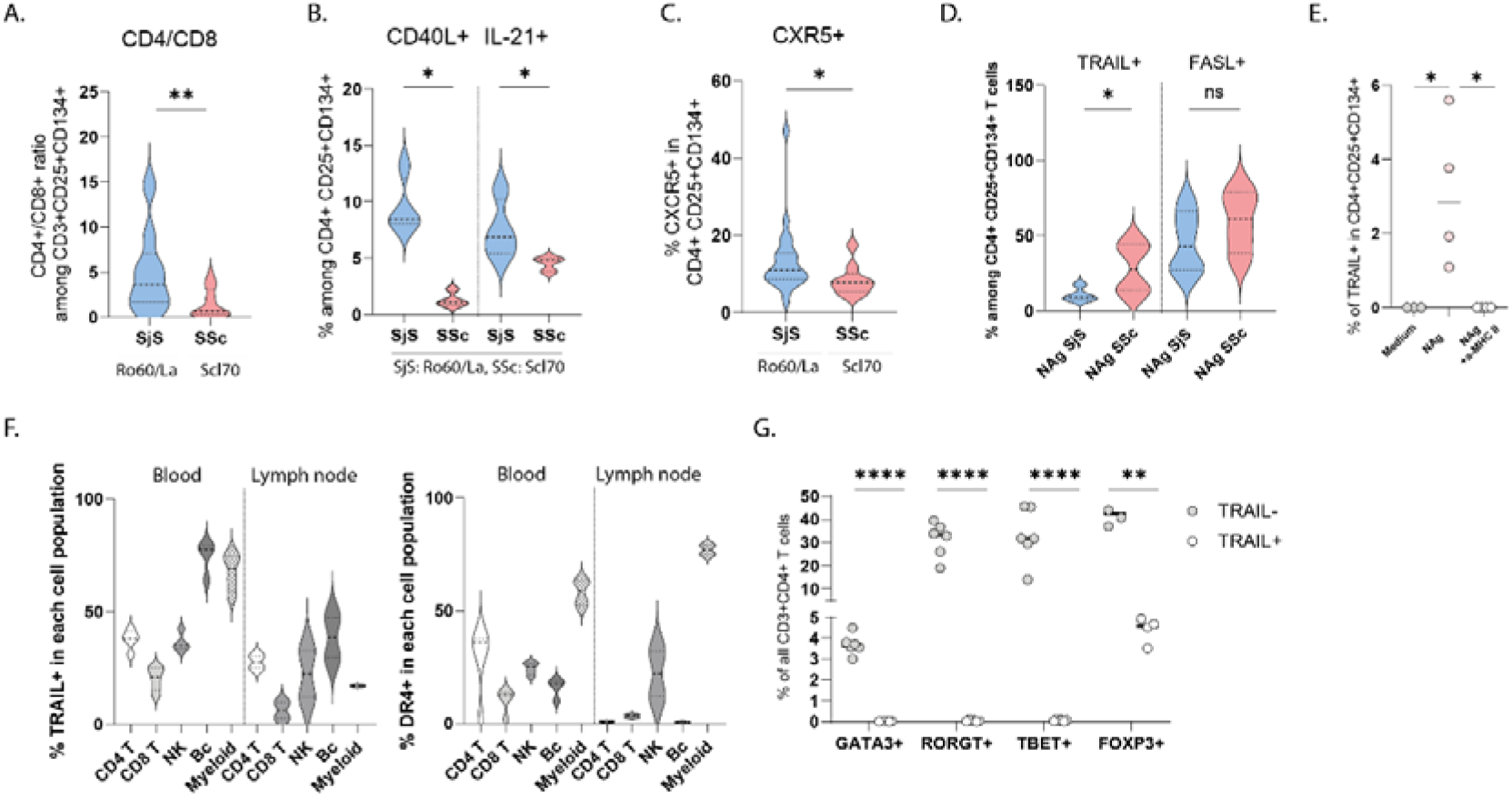
SjD blood features more antigen stimulus-responsive CD4+ T-cells, displaying signs of B-cell help, compared to fewer and TRAIL expressing antigen stimulus-responsive CD4+ T-cells in SSc. ArCD4+ T-cells, detected with the AIM assay as CD4+CD25+CD134+, were measured with flow cytometry on patients PBMCs that were stimulated with nuclear autoantigens (NAg) (Scl70 in SSc, Ro60/La in SjS). **(A)** Violin plots representing the ratio of arCD4+/CD8+ T-cells between SjS (n=37) and SSc (n=45). **(B)** Violin plots representing the proportion of arCD4+ T-cells expressing CD40L (n=6) and IL-21 (n=4) between SjS and SSc; **(C)** % of CXCR5 (Tfh-like) arCD4+ T cells; SjS (n=20) vs SSc (n=9). **(D)** of TRAIL and FASL between SjS (n=6) and SSc (n=6). (E) Proportion of arCD4+ T-cells (n=4) expressing TRAIL after 16-hour incubation with Ro60/La antigens and with/without MHC class II blockade. **(F)** Expression of TRAIL and TRAILR1/DR4 in blood/LN immune cells. Statistical differences: For (A-C) two-tailed unpaired t test with significance set at *p<0.05, **p<0.01, ****p<0.0001; **(D)** RM one-way ANOVA, with Dunnett’s multiple comparisons test, *p<0.05; (E) ordinary one-way ANOVA, *p<0.05.

### Potential functions of TRAIL+ CD4+ ISG T-cells

To gain more insight in the cells with which TRAIL+ CD4 T-cells might interact we analyzed expression of its receptor DR4 on T-cells and other immune cell populations. In blood DR4 was expressed by a proportion of all immune populations. In LNs it was predominantly expressed by NK cells and especially professional APCs *(<u>Figure 7F</u>)*. This pattern aligned with our immunohistochemical data indicating that TRAIL+ CD4+ T-cells seem to primarily interact directly with APCs. When cultured in vitro in the presence of anti-CD3/CD28 and IFNα, compared to other effector T-cell subsets, CD4+TRAIL+ ISG T-cells from SjD patients, SSc patients and healthy donors similarly displayed a reduced IFNγ production and increased expression of IL-10 and TRAIL, suggesting that TRAIL+ CD4+ ISG T-cells may exert regulatory rather than effector effects (Supplemental Figure 9A-C). To explore such a function, we isolated TRAIL+ CD4+ T-cells and conventional Tregs and compared their effects on effector CD4+ T-cells. Interestingly, TRAIL+ CD4+ T-cells suppressed CD4+ effector T-cell proliferation in similar fashion as conventional Tregs *(<u>Figure 8A, B</u>, <u>supplemental Figure 9D</u>)*. In line with this finding, co-culture experiments of antigen-stimulated monocytes, CD4+ T-cells and B-cells of SjD and SSc patients *(<u>Figure 8C</u>)* demonstrated that TRAIL blockade resulted in increased generation of IL-4, IL-17 and IFNγ producing T-cells *(<u>Figure 8D</u>)*. Furthermore, TRAIL/TRAIL receptor blockade resulted in increased formation of autoantigen-specific plasma cells *(<u>Figure 8E-GH</u>)* and production of ANAs *(<u>Figure 8I-K</u>)*. Taken together these exploratory analyses confirmed that prolonged antigen-stimulation assays can be used to study autoreactive T & B-cell responses and suggest that TRAIL+ CD4+ ISG T-cells and TRAIL restrict autoreactive responses in autoimmune CTDs.

**Figure 8.**
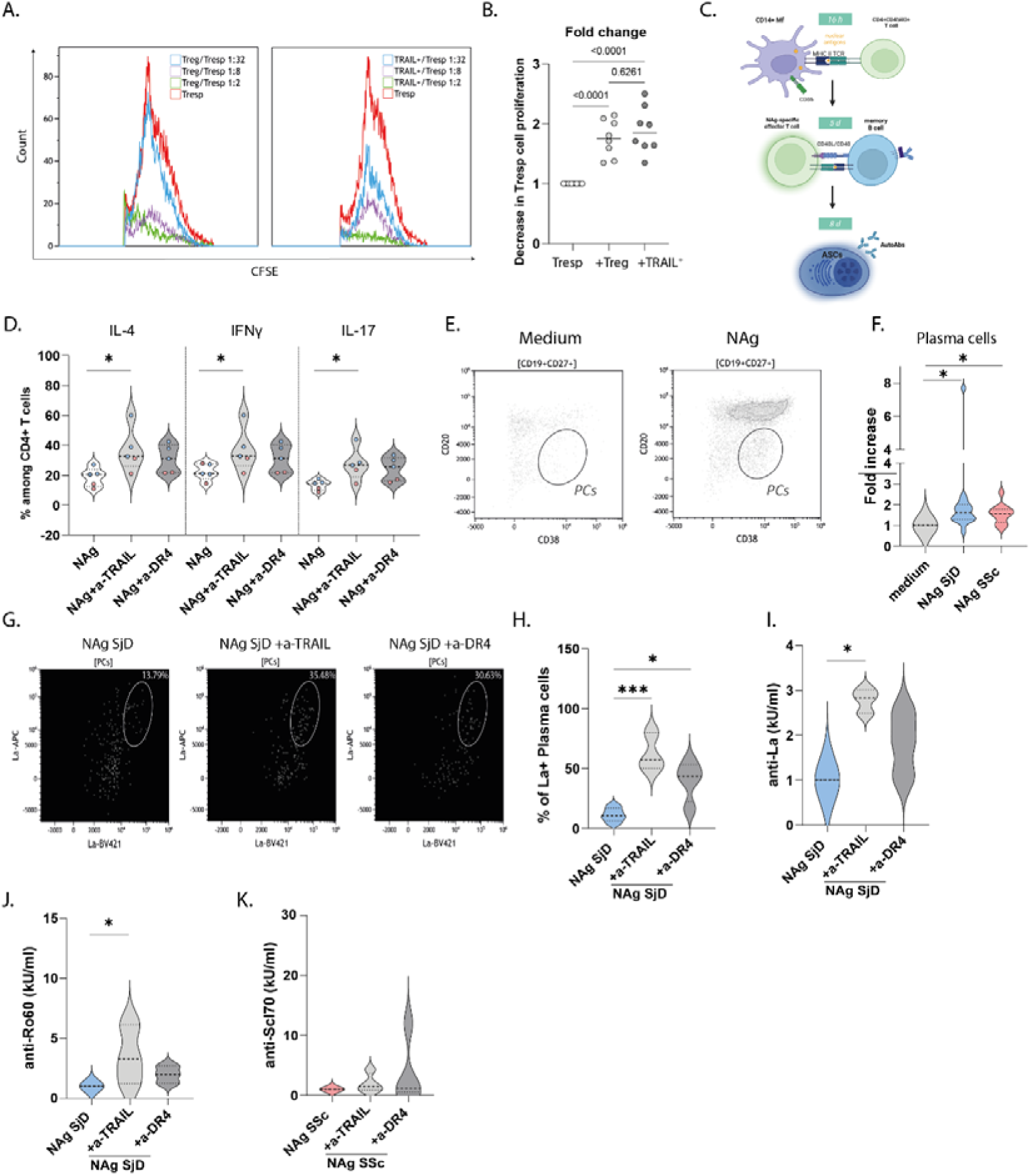
arCD4+ TRAIL+ ISG T-cells restrict plasma cell formation by suppressing APC mediated autoreactive humoral responses. **(A)** Overlay histogram flow cytometry plots illustrating proliferation of CD4+ effector responsive T-cells (Tresp) that were cultured alone or in the presence of either regulatory T-cells (Tregs) or TRAIL+ CD4+ T-cells at 1:2, 1:8 and 1:32 ratios (Treg/TRAIL+:Tresp). **(B)** Proliferation was measured with CFSE fluorescence emission in CFSE labeled Tresp cells and quantified as the fold change of the decrease in Tresp proliferation for all included donors (n=8). **(C)** Schematic representation of the in vitro mechanistic experiments unraveling the potential role of arCD4+ TRAIL+ T-cells towards modulating APC/T/B cell interactions. Co-cultured CD4+ memory T-cells, B-cells and monocytes were stimulated with NAgs (n=3 Ro60/La and n=2 Scl70) in presence or not of anti-TRAIL/anti-DR4 for 5 to 8 days. **(D)** Proportion of CD4+ memory T-cells expressing IL-4, IFNγ, IL-17. **(E)** Representative plots (n=10) illustrating in vitro generation of plasma cells that are further quantified in **(F)** as fold change increase on % of CD19+CD27+CD20^low^CD38+ plasma cells/all live cells. **(G)** Flow cytometry plots of one representative experiment (n=4) showing detection of La-specific plasma cells in the absence/presence of TRAIL/TRAILR blockade. **(H)** Percentage of La-specific plasma cells among live plasma cells (n=4). (I-K) Quantification of autoantibody (anti-La/Ro60/Scl70) production in cell culture supernatants (n=6 per condition). Statistics: in (B) RM one-way ANOVA, with Dunnett’s multiple comparisons test, *p<0.05; in (D), (F) non-parametric Kruskal-Wallis test, *p<0.05; in (B), (H), (I), (J), (K) ordinary one-way ANOVA, with Tukey’s multiple comparisons test, *p<0.05, ***p<0.001.

## Discussion

Autoreactive CD4+ T-cell responses are considered to orchestrate autoimmune pathology in CTDs (26). The efficacy of autologous stem cell transplantation and anti-CD19 CAR T-cell therapy in treating CTDs highlights the need to eliminate autoreactive cells (27, 28). However, poor understanding of mechanisms of immunological disease activity in CTDs hinder the development of safer, more personalized tolerogenic therapies. Here, we demonstrate that strong differences in CD4+ T-cell infiltration in disease affected tissues of individuals with CTDs correlate to differences in CD4+ T-cell activation and regulation within active locoregional LNs and to circulating arCD4+ T-cells in blood.

Phenotypic and functional characterization of pathogenic T-cell responses in affected tissues and active LNs has been challenging due to difficulties in identifying and isolating these rare cell populations. Therefore, T-cells have been primarily studied in patients’ blood which limited the insights on the systemic and local function of these cells. Therefore we analyzed if [^18^F] FLT PET scans allow for detection of active locoregional LNs and affected tissues, enriched for disease-involved T-cells. We found that the [^18^F] FLT-positive LNs contained a higher number of proliferating immune cells compared to the [^18^F] FLT-negative ones, with enhanced formation of GC-like structures and clonal expansion of T-cells and B-cells. These findings align with previous studies in patients with melanoma and head and neck carcinomas (20, 29) suggesting that [^18^F] FLT PET scan may effectively sample active LNs enriched with disease-involved proliferating immune cells.

Combining [^18^F] FLT-PET scans with multiplex immunohistochemistry, single-cell RNA sequencing, and T-cell activation assays, we found that SjD patients exhibit robust APC, B-cell, and T-cell interactions forming germinal center-like structures in active LNs, associated with a pleiotropic CD4+ T-cell effector response and extensive APC/CD4+ T-cell/B-cell aggregates in inflamed SGs. In contrast, SSc active lymph nodes showed limited APC/T/B-cell interactions and an expanded population of extrafollicular CD4+ TRAIL+ ISG T-cells. This qualitative difference in the CD4+ T-cell response may be explained by either a stronger innate immune response in SjD, which could trigger presentation of autoantigens together with stronger pro-inflammatory signals, or by the inherent properties of the autoantigens themselves (30)—particularly Ro/SSA and La/SSB, which are highly immunogenic (31) and widely expressed in epithelial tissues. Such factors may drive persistent B-cell activation and promote the formation of tertiary lymphoid structures (32). In contrast, SSc patients displayed a modestly increased arCD4+ Treg/effector cell ratio in active LNs and blood, consistent with previous findings of elevated Tregs in SSc blood with uncertain suppressive function (33, 34).

These results also align with studies in NOD mice showing that regulatory T-cell activity influences development of follicular versus extrafollicular humoral responses (35, 36) and with our earlier findings in 135 SSc patients of increased skin-infiltrating CD8+ T-cells but not CD4+ T-cells (9).

The AIM assay allowed for enrichment and analysis of functional markers on arCD4+ T-cells from blood that displayed similar polarization as CD4+ T-cells from active LNs and that were part of clones that were expanded in active LNs and disease-affected tissues. Although the AIM assay is commonly used to identify antigen-reactive CD4⁺ T cells, they have inherent limitations for this purpose.

Despite our validation experiments, i.e. HLA-DR blocking and an autoreactive peptide-MHC class 2-tetramer, which showed a relatively large overlap between arCD4+ T-cells and tetramer positive T-cells, the response of a large fraction of peripheral blood CD4+ T-cells to especially the combination of Ro60 and La stimulation suggests a response to stimulation that is broader than only involving antigen-specific cells. Insight into the antigen-specificity of arCD4+ T-cells requires recombinant expression of the full length T-cell receptors of these cells which is beyond the scope of the current study. Despite this limitation, we found that this assay highly enriched for expanded tissue residential CD4+ T-cells. Tetramer staining of available autoreactive epitopes in a number of MHC matched subjects showed that these included a significant proportion of autoreactive T-cells. Even after pre-selection of active LNs with [^18^F] FLT-PET only 7% of CD4+ T-cells in active LNs was clonally expanded. However, AIM responsive CD4+ T-cells from blood were circulating clonal counterparts of CD4+ T-cells from active LNs and disease-affected tissues of which 35% were clonally expanded, including the most expanded tissue residential clones. As such this validates antigen-stimulation based assays for experiments with circulating CD4 T-cells enriched for disease-involved clones in CTDs. This was confirmed by our finding of similar activation and polarization of circulating arCD4+ and tissue residential CD4+ T-cells. As further validation of the use of ex vivo antigen-stimulation assays of peripheral blood mononuclear cells we found that a monocyte-T-cell-B-cell co-culture can be used to study the formation of effector CD4+ T-cells, autoreactive plasma cells and autoantibodies.

Active LNs of especially SSc patients contained a uniquely expanded population of TRAIL+ CD4+ ISG T-cells. TRAIL is a type II transmembrane protein belonging to the TNF superfamily, closely resembling the FAS ligand. TRAIL triggers cell apoptosis by binding to its death receptors (DR4/DR5), which activates apoptosis signaling via the caspase cascade. However, primary immune cells are often resistant to TRAIL-induced death (37) by concurrent signaling mechanisms, such as CD40-CD40L, or by expressing decoy receptors (38). The functional role of TRAIL, beyond its potential to kill cancer cells, is not well defined. CD4+ TRAIL+ T-cells have been primarily analyzed in peripheral blood and affected tissues with vasculopathy and atherosclerosis, where they were found to contribute to disease activity by killing vascular smooth muscle cells (21–23). In these studies, no ISG gene signature in CD4+TRAIL+ T-cells was reported. In contrast, the unique and novel T-cell subset that we observed and termed CD4+ TRAIL+ ISG T-cells was primarily detected in the active LNs of SSc patients and rarely in the blood or affected tissues of patients with SSc and SjD. This subset was characterized by the combined expression of the death receptor TRAIL and a type I IFN gene signature specific to this cluster of cells. This novel CD4+ TRAIL+ ISG T-cell subset exhibits a clonally expanded, naïve-like memory phenotype with mixed features of quiescence and activation. Of interest, CD4+ ISG T-cells exhibiting antigen specificity and TCR expansion were recently described in patients with systemic lupus erythematosus (39). The ISG response program in these T-cells might reflect recent interferon mediated activation of activation-prone T-cells. Of interest, in active LNs of SSc patients this did not translate in generation of canonical CD4+ T-helper cells. Instead, trajectory analysis indicated a separate pathway of development for these cells that separated from development of T-helper or regulatory CD4+ T-cells.

On the one hand TRAIL+ CD4+ ISG T-cells may involve bystander T-cells, albeit that their relative strong clonal expansion argues against a bystander activated nature. Alternatively, they may involve an autoreactive T-cell subset with a regulatory function that counteracts proinflammatory autoreactive responses. This was suggested by preliminary experiments showing that TRAIL+ T-cells from healthy persons may suppress CD4+ effector T-cells and that in CTDs TRAIL blockade enhanced the autoreactive effector response in an ex vivo antigen-stimulation monocyte-T-cell-B-cell co-culture.

SSc may feature a relative increase in these cells because of a low immunogenic autoantigen response, e.g. lacking CD40L stimulating signals, or because of a different polarization of the innate immune activation. Such mechanisms are suggested by observations that SjD features a broad range of antibody specificities (40), whereas SSc involves more limited antibody diversity and lower titers (41). In previous clinical studies we observed predominant activation of CD8+ over CD4+ T-cells in disease-affected tissues of patients with early severe SSc. In contrast, we observed increased CD4+ helper T-cells in the peripheral blood of patients with a longstanding and complicated disease course (42). This was correlated to a strong reduction in soluble TRAIL levels in these patients with a long-standing and complicated disease (data not shown)). Collectively, these findings suggest that TRAIL might protect against adaptive immune activation by restricting autoreactive T-cell and B-cell mediated disease severity. Taken together, TRAIL+ CD4+ ISG T-cells were relatively enriched in SSc patients. Bona fide studies in in vivo models will be required to proof if TRAIL+ CD4+ ISG T-cells concern bystander activated cells or a regulatory subset.

### Future perspectives

The study cohort of matched blood, LN and affected tissue biopsies consisted of patients with early, severe and progressive disease. Studies in prospective cohorts in patients from the full CTD spectrum are required to study correlations between CD4+ T-cell polarization, clinical disease manifestations and course. Experimental studies are required to answer open questions: why are CD4+ TRAIL+ ISG T-cells highly present in SSc active LNs but not in SjD? Are they regulated by the type of autoantigen (Ro/La bind RNA, while Scl70 binds DNA) or at the antigen-presentation level? Can we use these findings to design novel therapeutic approaches to tackle systemic autoimmune diseases, such as including adjuvants in tolerogenic vaccines that may divert the differentiation of T-cells towards an immunoregulatory phenotype? Such novel therapies might benefit a broader spectrum of autoimmune diseases, including diseases such as systemic lupus erythematosus, rheumatoid arthritis, and multiple sclerosis.

## Methods

### Sex as a biological variable

Our study examined both male and female patients and similar findings are reported for both sexes.

### Study participants

All SSc and SjD patients (aged >18) that donated whole blood (n=45 SSc and n=37 SjD) and/or LN (n=4 each disease), salivary gland (parotid) (n=3 SjD), skin (forearm) (n=2 SSc) biopsies, were diagnosed with established disease according to the ACR EULAR 2014 classification criteria (43) and European American consensus group classification criteria (44) respectively. SSc and SjD patients with overlapping syndromes were excluded from the study. SjD patients seropositive for anti-Ro60 and/or anti-La antibodies were included, while all SSc patients were seropositive for anti-Scl70 (anti-topoisomerase I). Diagnosis of early diffuse cutaneous SSc was performed according to the VEDOSS criteria (45) and presence of anti-Scl70 antibodies, a disease duration (from first non-Raynaud symptom) of < 3 years and progressive disease in the past year, as defined by either 1. an increase in mean Rodnan skin score (mRSS) > 10 points / > 25%, or 2. a decrease in forced vital lung capacity > 10%, because of an increase in interstitial lung disease. From all patients that donated tissue biopsies, paired peripheral blood mononuclear cells (PBMCs) were also available. Clinical characteristics for all patients are provided in Supplemental Table 1. Blood samples from healthy volunteers (n=30) were collected from Sanquin blood bank, Nijmegen, the Netherlands (project number: NVT 0397-02) from individuals that consented to donating blood for medical research.

### PET/CT acquisition and analysis

All patients were instructed to drink 1 L of water before imaging and received 10 mg furosemide intravenously to stimulate urinary tracer excretion. An integrated PET-CT scanner (Biograph mCT, Siemens, Knoxville, TN, USA) was used for data acquisition. Emission images were acquired one hour after intravenous injection of 250±10% MBq of [^18^F] FLT (Cyclotron B.V., VU Medical Centre, Amsterdam, The Netherlands). The images were corrected for attenuation using low-dose CT and reconstructed using the ordered-subsets expectation maximization (OSEM) algorithm, following EANM guidelines [DOI 10.1007/s00259-014-2961-x]. Low-dose CT scan was used for anatomical correlation.

The PET/CT image sets were assessed by a certified nuclear medicine physician for the presence or absence of [^18^F] FLT uptake in LNs. Three-dimensional regions of interest were placed manually over every LN by using a CE-marked viewing software Agfa Enterprise on multiple slices. Maximum and mean standardized uptake values (SUV_max_, SUV_mean_) were derived for LNs of interest. LNs were annotated as negative when uptake of the tracer was not higher compared to the surrounding connective tissue and positive for uptake that was higher than the cutoff defined in preceding studies (19, 20).

### Sample preparation

PBMCs were isolated from whole blood by using Ficoll-Paque PLUS (Sigma Aldrich, GE17-1440-03) density centrifugation and were cryopreserved as previously described (42) and stored in liquid nitrogen until future use. After thawing and washing, PBMCs where cultured in complete RPMI 1640 medium with GlutaMAX™ (Gibco, ref 72400-021), supplemented with 100 IU/ml penicillin, 100 mg/ml streptomycin, 100 mg/L sodium pyruvate, and 10% human pooled serum (HPS). LN biopsies (three-five 1.25 mm biopsies from each donor) obtained with the use of HistoCore autobiopsy system (cat # HC18100) were rinsed in complete RPMI medium in 6-well plates and with the help of a syringe staple passed through a 70 µm cell strainer to obtain single-cell suspensions. After washing with PBS, red blood cells were lysed using ice-cold erythrocyte lysis buffer (155 mM NH_4_Cl, 12 mM KHCO_3_, 0.1 mM EDTA in PBS) for 2 minutes at room temperature. For salivary gland and skin tissue disaggregation, tissue fragments were minced with a scalpel and enzymatically digested by using 0.1 mg/ml DNAse I (DN-25, Merck, Darmstadt, Germany) and 0.1 mg/ml Liberase TM (5401127001, Roche, Vienna, Austria) in plain RPMI 1640 for 60 minutes at 37 °C on a roller bank. For skin, 3-mm punch biopsies’ mechanical dissociation was additionally used before and after enzymatic dissociation using a gentle MACs dissociator (program h_skin_01). The digested fragments were passed through a 70 µm cell strainer to obtain a single-cell solution and washed with complete RPMI medium.

### Sample preparation for single-cell RNA sequencing

For single-cell RNA sequencing experiments utilizing tissue biopsies and PBMCs, single-cell suspensions were washed twice with PBS before they were stained with Fc block (10 minutes at 4 °C; BD Biosciences, cat 564219), and then with CITE-seq antibody cocktail. For CITE-seq, cells were stained with the TotalSeq™-C Human Universal Cocktail, V1.0 (Biolegend, cat#399905) according to the manufacturer’s instructions. Then, viable single-cell suspensions were FACs sorted (Supplemental Figure 10A) in a BD FACSMelody™ Cell Sorter in cooled sorting tubes (4 °C) and immediately processed according to manufacturer’s guidelines (10x Genomics) for Chromium Single Cell Immune Profiling using Chromium Next GEM Single-cell 5’ HT Reagent Kits (v.2) and the recommended reagents, supplies and equipment. Sorted cells were loaded into the 10x chromium cell controller at a density of 1200 cells/µl to optimally capture up to 10,000 cells per sample. Quality control of the generated libraries was performed using the Qubit 1x dsDNA HS assay kit (Invitrogen) and the Bioanalyzer (Agilent). The sequencing of the barcoded cDNAs was performed on the Nextseq500 (Illumina) using paired end reads. The average sequencing depth was at least higher than 70K raw reads per cell.

### Multimodal single-cell RNA sequencing analysis

Raw data were prepared for analysis using the Cell Ranger (v.7.2.0; 10x genomics) and FASTQ reads were aligned from gene expression (GEX), ADT/HTO and V(D)J sequencing libraries to the prebuilt GRCh38 human transcriptome. TRA and TRB sequences were annotated using the Cell Ranger VDJ function from 10x Genomics. Preprocessing of the raw data and subsequent analysis was performed in R (version 4.4.0) using the Seurat (5.1.0) package (46). Quality control measures of the count matrix were applied to filter out T-cells with a mitochondrial gene content exceeding 5% and cells with too low or too high feature counts per cell (customized for each dataset). Following this, CD3+ T-cells were computationally isolated and sorted for separate analyses based on unsupervised clustering and expression of CD3E/D/G. CD4+ T-cell clustering was then performed based on gene expression of CD4 measured by Seurat and CD4+ T-cell clusters were identified based on each cluster’s differentially expressed genes (DEGs) and differentially expressed proteins (DEPs; a CITE-seq panel of 137 antibodies was used). Cell type annotation was further validated based on the expression (gene and protein) of canonical subset markers and with the use of joint density (47) to identify co-expressed genes relevant to certain T helper clusters. To ensure high purity and exclude CD4, CD8 double positive T-cells, we afterwards filtered out T-cells that had CD8A normalized mRNA expression levels greater than 1 (padj < 0.01 and log2FC >0.5).

For primary dimensionality reduction, non-negative matrix factorization (NMF) was utilized, followed by Uniform Manifold Approximation and Projection (UMAP) and Louvain clustering using top 40 NMF components, as described by Singh et al. (48). Counts of the CITE-seq antibodies were normalized with the use of the centered log ratio transformed (CLR) counts. To identify DEGs and differentially DEPs within each transcriptionally distinct cluster, the *FindAllMarkers* function (with logistic regression; LR as testing method) in Seurat was employed. These DEGs and DEPs were annotated based on their characteristics, and the R package *pheatmap* (Kolde, R. (2019). *pheatmap*: *Pretty Heatmaps* (R package version 1.0.12) was used to visualize them across multiple cell types. Top DEGs and DEPs per cluster were statistically defined using LR test and the Benjamini-Hochberg method to adjust for multiple testing.

### Integration of T-cell receptor clonotype analysis with single-cell RNA sequencing data

T-cell receptors (TCR)s were annotated using the Cell Ranger VDJ pipeline. The *scRepertoire* R package (v1.153) was utilized to identify and analyze TCR clonotypes based on TCR alpha and beta chains as well as CDR3 sequences. Clonotype data were integrated with Seurat to generate gene expression and cluster information using the *combineExpression* function. Cells sharing the same paired TCRαβ genes were annotated as belonging to the same clonal lineage. To visualize shared clones across clusters or tissues and to determine expanded TCR clonotypes in paired blood, LNs and affected tissue biopsies, a chord diagram was generated using the *getCirclize* function from the R package *circlize*. Heatmaps were created using the *ComplexHeatmap* R package (v2.13.153,55). Additionally, shared expanded TCR clonotypes in paired blood, LNs and affected tissue biopsies were visualized using the *circlize* R package. All scripts used for analysis of the bioinformatic data of this manuscript can be found here; https://github.com/PrashINRA/TRAIL_Manuscript.

### Analysis of tissue arCD4+ TCR clonotypes paired with single-cell RNA sequencing data

To analyze TCR clonotypes shared between blood, LNs and affected tissues, we used paired single-cell RNA and TCR sequencing in all three compartments of SjD and SSc patients (blood, LNs, affected tissues). T-cells that had at least one annotated α and one annotated β chain in the TCR data were classified as matching if a cell in the paired tissue/blood data exhibited the exact same α and β chain composition. Only T-cells with at least one annotated α and one β chain were included in analyses comparing matching blood, LN and affected tissue cells. Two cells were considered part of the same T-cell clone if they shared both the exact same α and β chains, as determined by the amino acid sequence. If cells possessed multiple α and β chains, they were deemed matching only if all detected α and β chains were identical. This strict definition was applied to ensure that each pair of cells within the same clone exhibited complete similarity in their detected TCR chains, indicating with high probability that they originated from the same T-cell clone. TCR data was also used to quantify clonal expansion by counting the number of cells in each clonotype. Detection and characterization of arCD4+ T-cells in the LNs and affected tissues were achieved through single-cell RNA sequencing, CITE-seq, and TCR sequencing across blood, LNs and affected tissues. TCR sequences derived from ex vivo blood arCD4+ T-cells, that were FACs sorted and subjected to single-cell RNA sequencing analysis, served as unique identifiers, akin to molecular barcodes, enabling the mapping of identical TCR sequences across blood, LNs, and affected tissues.

### Flow cytometry and intracellular staining

0.5-1 x 10^6^ PBMCs/single-cell suspensions were first labeled with ViaKrome 808 fixable viability dye (1.5:1000 in PBS) for 30 minutes at 4 °C to exclude dead cells, followed by staining with fluorescently labeled extracellular antibodies (Supplemental Table 2) for 20 minutes at room temperature. For intracellular antibody staining (Supplemental Table 3), cells were fixed and permeabilized using the Cyto-Fast™ Fix/Perm Buffer Set (Biolegend) according to the manufacturer’s guidelines. To facilitate detection of intracellular cytokines, cells were pre-stimulated with 12.5 ng/ml phorbol 12-myristate 13-acetate (PMA) (Sigma), 500 ng/ml ionomycin (Merck), and 5 µg/ml brefeldin A (Merck) before staining. Samples were acquired on a Beckman Coulter Cytoflex LX 21-color flow cytometer immediately after staining.

To detect antigen-specific T-cells and B-cells with the use of fluorescent tetramers (Supplemental Table 4) the flow cytometric protocol was as described earlier with the addition that after cell viability staining, cells were incubated with designated fluorescent tetramers at room temperature in the dark for 30 minutes before the staining for extracellular markers was performed. Tetramers to detect Ro60-specific T-cells were obtained through the NIH Tetramer Core Facility (contract number 75N93020D00005). Streptavidin labeled tetramers to detect autoreactive La-specific B-cells were kindly provided by Dr. Mathijs Broeren.

### AIM assay

Cryopreserved PBMCs were thawed and resuspended in RPMI 1640 with 10% HPS medium and seeded in 96-well u bottom plates with 250,000 cells per well (4-plo per condition). PBMCs were cultured for 16 hours at 37 °C and 5% CO_2_ in the presence of negative control (DMSO vehicle), positive control (1 µg/ml) Staphylococcal enterotoxin B (SEB) (Sigma) and recombinant human proteins. Experimental replicates (quadruplicates) were conducted for all conditions. The human recombinant proteins comprised influenza A H1N1 nucleoprotein (NP, Sino Biological, 11675-V08B) at 2 µg/ml and La/SS-B ribonucleoprotein (La, Prospec, PRO-327), 60 kDa SS-A/Ro ribonucleoprotein (Ro60, Prospec, PRO-329), DNA topoisomerase 1 (Scl70, Prospec, ENZ-306) all at a concentration of 1 µg/ml. After a 16-hour incubation, cells were washed with PBS and used for flow cytometry staining. The gates for arCD4+ T-cells were drawn based on increased expression of the early T-cell activation markers CD25/CD134 (OX40) as response to (auto)antigen stimulation (representative plots and gating strategy are provided in Figure 4B and Supplemental Figure 10B). In experiments where the assay was combined with intracellular staining, after the 16-hour incubation, 5 µg/ml Brefeldin A (Merck) was added, and cells were cultured for additional 4 hours to facilitate detection of intracellular proteins. To block the interaction of HLA class II complex and CD4+ T-cells, 1 µg/ml of purified anti-human HLA-DR, DP, DQ (clone; Tü39, Biolegend, 361702) antibody was added simultaneously with antigen stimulation.

arCD4+ T-cells that were responsive to the anti-nuclear antigens La, Ro60 and Scl70 were isolated using FACS (see for gating strategy Supplemental Figure 10B) and processed for single-cell RNA/TCR sequencing to analyze arCD4+ T-cell responses and generate arCD4+ TCR repertoire libraries. Single-cell RNA sequencing data analysis of anti-nuclear antigen stimulated T-cells was performed with the same workflow that was previously described.

### Immunohistological analysis

LN, salivary gland and skin biopsies were formalin fixed, paraffin embedded (FFPE), sectioned (5.0 µm thickness) and subsequently stained with hematoxylin and eosin (HE). For immunohistochemistry, the FFPE tissue sections were deparaffinized with xylol and rehydrated with ethanol. Antigen retrieval was performed in a 10 mM sodium citrate buffer (pH 6.0) in water either at room temperature or by heating the slides for 30 minutes at 97° C. Blocking of endogenous peroxidase was conducted with the use of 3% H_2_O_2_ in PBS. Primary (Supplemental Table 5) and appropriate secondary (BrightVision Poly-HRP, Immunologic DPVO55HRP or Envision Flex HRP, DAKO) antibody labeling was performed with 3’3’-diaminobenzene (bright DAB, Immunologic or DAKO) reagent and sections were counterstained with hematoxylin. For multiplex immunofluorescent staining, slides were stained using an automated platform with the Opal 7-color Automation IHC kit (NEL801001KT; PerkinElmer) on the BOND RX IHC & ISH Research platform (Leica Biosystems), following previously described protocols (9). We refer to Supplemental Table 6 for a full list of the antibodies that were used. Slides were then scanned using the Automated Quantitative Pathology Imaging System (Vectra V.3.0.4, PerkinElmer) at 4x magnification for an overview. Multispectral images of tissue biopsies were annotated with Phenochart (V.1.0.9, PerkinElmer) and scanned at 20x magnification. Spectral unmixing of the Opal fluorophores was performed using

InForm software (V.2.4.2, PerkinElmer), and the multichannel images were digitally merged. For quantitative analysis, digital scans of whole tissue biopsies (three sections per biopsy per donor) were quantified using QuPath-0.4.4 (50).

### Co-culture suppression assays

The suppressive capacity of FACs sorted (see Supplemental Figure 9D for the gating strategy) CD4+CD25highTRAIL-Tregs and CD4+TRAIL+CD25-T-cells was assessed using co-culture suppression assays. Tregs and TRAIL+ CD4+ T cells were sorted from fresh blood PBMCs after CD4+ T-cell enrichment was performed with a MojoSort™ Human CD4 T Cell Isolation Kit (Biolegend, cat#480010) according to manufacturer’s instructions and followingly co-cultured with CFSE-labeled CD4⁺CD25⁻TRAIL-responder T cells (Tresp) at varying ratios, in the presence of anti-CD3/anti-CD28 mAb-coated beads (ThermoFischer, cat#11131D), bead-to-cell ratio 1:5, for 3 days. Tresp proliferation was quantified by CFSE dilution, as previously described (51).

### B-cell help immunoassay

To evaluate B-cell help function of arCD4+ T-cells, we utilized a sensitive immunoassay, developed by Ansari et al. (52). In this assay, CD19+ B-cells, CD4+CD45RO+ memory T-cells and CD14+ monocytes were FACS sorted (Supplemental Figure 10C) from SSc and SjD patients’ blood and co-cultured at 1:1:0.5 ratio with or without stimulation with recombinant human La, Ro60, Scl70 for 8 days at 37 °C and 5% CO2. Generation of (autoreactive) plasma cells was assessed with flow cytometry and plasma cells were gated as CD19+CD27+CD38++CD20low (Supplemental Figure 11). For detection of La-specific B-cells, La-APC/BV421 tetramers were used in the flow cytometric extracellular staining. To block TRAIL-TRAIL Receptor pathway signaling, recombinant human anti-TRAIL (Biolegend, 308202) and anti-TRAILR1/CD261 (DR4; Biolegend, 307201) antibodies were used at 10 µg/ml.

### Quantification of anti-nuclear autoantibodies

To measure Ro60, La and Scl70-specific IgG antibodies in the co-culture supernatants of the B-cell help immunoassay experiments, an EliA Symphony assay (Thermo Fisher Scientific, Inc., Waltham, MA) was used as previously described (53). The analyses were conducted using an immunoassay analyzer (ImmunoCAP 250, Thermo Fisher Scientific, Inc.). Values are presented in kU/ml and were corrected based on the number of live B-cells per sample.

### RNA isolation and quantitative real-time polymerase chain reaction

RNA isolation was carried out using RNAeasy (Qiagen) following the manufacturer’s instructions. The RNA concentration was then measured with a Nanodrop spectrophotometer (Thermo Scientific, Waltham, MA, USA), and any genomic DNA was removed using DNase I. Up to 1 μg of RNA was reverse-transcribed into cDNA in a single-step reverse transcription PCR at 39°C using an oligo dT primer and 200U M-MLV reverse transcriptase (All Life Technologies) in a thermocycler. Gene expression in the resulting cDNA was measured using 0.2 mM validated primers (Biolegio, Nijmegen, the Netherlands; see Supplemental Table 7) and SYBR Green master mix (Applied Biosystems, Waltham, MA, USA) in a quantitative real-time polymerase chain reaction (qPCR). Relative gene expression (-ΔCt) was calculated based on the average expression of three reference genes: *GAPDH, TBP* and *RPS27A*.

### Computational analysis of flow cytometry data

For dimensionality reduction of flow cytometry data, viSNE analyses were conducted using the web-based analysis software Cytobank (http://cytobank.org/) (54). Initially, each file was gated on live CD3+ T-cells and then on certain T-cell populations of interest for each analysis. Subsampling followed the visNE algorithm, analyzing 14,060 events per sample. The t-SNE axes of the viSNE maps were generated using the markers CD3, CD4, CD8, CCR4, CD25, CD69, CXCR5, CD28, CCR4, CCR6, CXCR3, CD45RA, CD27, PD-1, ICOS, CD154.

The levels of each protein marker were normalized to the maximum value observed for the respective channel within each sample. For unsupervised analysis, T lymphocyte populations of interest were manually gated and isolated using the Beckman Coulter plug-in of Kaluza (v2.1.2) software. The files were categorized into comparison groups based on patient origin and T-cell population. Unsupervised clustering of lymphocyte and T-cell frequencies was performed with the CITRUS tool, using abundance-based clustering, a minimum cluster size of 2% of total cells, and a false discovery rate of 1%. Clustering was based on the expression levels of cell surface and intracellular markers from the viSNE analysis. The Citrus "cluster tree" illustrated the clustering hierarchy, with nodes scaled according to cell frequency in each cluster. Clustering was evaluated using prediction analysis for microarrays (PAM) to identify features that distinguish certain T-cell clusters with each other and/or certain T-cell clusters between patients with SjD and SSc.

### Statistical analysis

Data visualization and statistical comparisons between experimental groups were carried out using R Studio (version 4.1.3) and Prism software (GraphPad 9.0.0, San Diego, CA, USA). Data are presented as means ± SEM. Two-tailed unpaired Student’s T-tests were used for comparisons between two groups, while one-way or two-way ANOVA with Tukey’s multiple comparisons test or non-parametric Kruskal Wallis test was applied for comparisons among multiple groups. Spearman’s correlation was used to analyze relationships between two variables that were not normally distributed. P values less than 0.05 were considered significant. To compare significance in cell frequency of single-cell RNA sequencing clusters the Wilcoxon test, corrected for multiple comparisons, was used. The specific statistical tests used for each analysis or experiment are detailed in the figure legends.

### Study approval

This study was approved by the local research ethics committee of Radboud University Medical Center, the Netherlands (study number(s): NL 67672.091.18) and all participants provided signed informed consent according to the principles of the Declaration of Helsinki. All patient related procedures were executed in accordance with the relevant Dutch legislation and reviewed by an accredited research ethics committee.

### Materials availability

This study did not generate new unique reagents. Data and code availability: The datasets and scripts used in this manuscript can be found here; https://github.com/PrashINRA/TRAIL_Manuscript. This study did not generate any unique code.

## Lead contact

Further information and requests should be addressed to the lead contact, Dr. Rogier Thurlings.

## Author contributions

T.I.P, H.K and R.M.T conceptualized the project. T.I.P executed the wet lab experiments and performed their data analysis. Technical support was provided by B.W, L.J.A.W and E.V. Bioinformatic analysis of single-cell RNA sequencing data was performed by P.S. P.K assisted with bioinformatic analyses at the revision process of the manuscript. Funding WAS acquired by R.M.T. This study investigation was carried out by T.I.P, A.C, H.K, I.J.M.V, M.A.H and R.M.T. Methodology was developed by T.I.P, P.S, A.C, R.L.S, K.W.M, I.J.M.V, E.A, M.A.H and R.M.T. Supervision was provided by A.C, X.H, H.K, P.K, R.L.S, E.A, I.J.M.V, M.A.H and R.M.T. Visualization was performed by T.I.P, K.M.H and E.A; The original draft was written by T.I.P and R.M.T, and the manuscript was reviewed and edited by T.I.P, A.C, X.H, H.K, K.W.M, M.V, E.A, I.J.M.V, M.A.H and R.M.T.

## Declaration of generative AI and AI-assisted technologies

No generative AI nor AI-assisted technologies were used in the writing process.

## Supplemental material

Supplemental Figures 1-14

Supplemental Tables 1-7

## Acknowledgements

We acknowledge a. the work performed by our student Myrthe van der Waal in the optimization of the in vitro functional assays during her internship in our department of Rheumatology b. the work of Dr. Mark Gorris and Kiek Verrijp in setting up multiplex immunofluorescence panels. We thank Dr. Massis Krekorian for his assistance in the collection and processing of patients’ biopsies for single-cell RNA sequencing, Annika Decker for her assistance in experiment evaluating antigen-presentation in AIM assay. Finally, we thank Pim Kloosterman for his bioinformatic support.

