## Supplemental materials for "Imaging guided single-cell multiomics unveils shared autoreactive CD4+ T-cell responses in blood, locoregional lymph node and affected tissues of patients with systemic autoimmunity"

### Supplementary Figures

**Supplementary Figure 1**

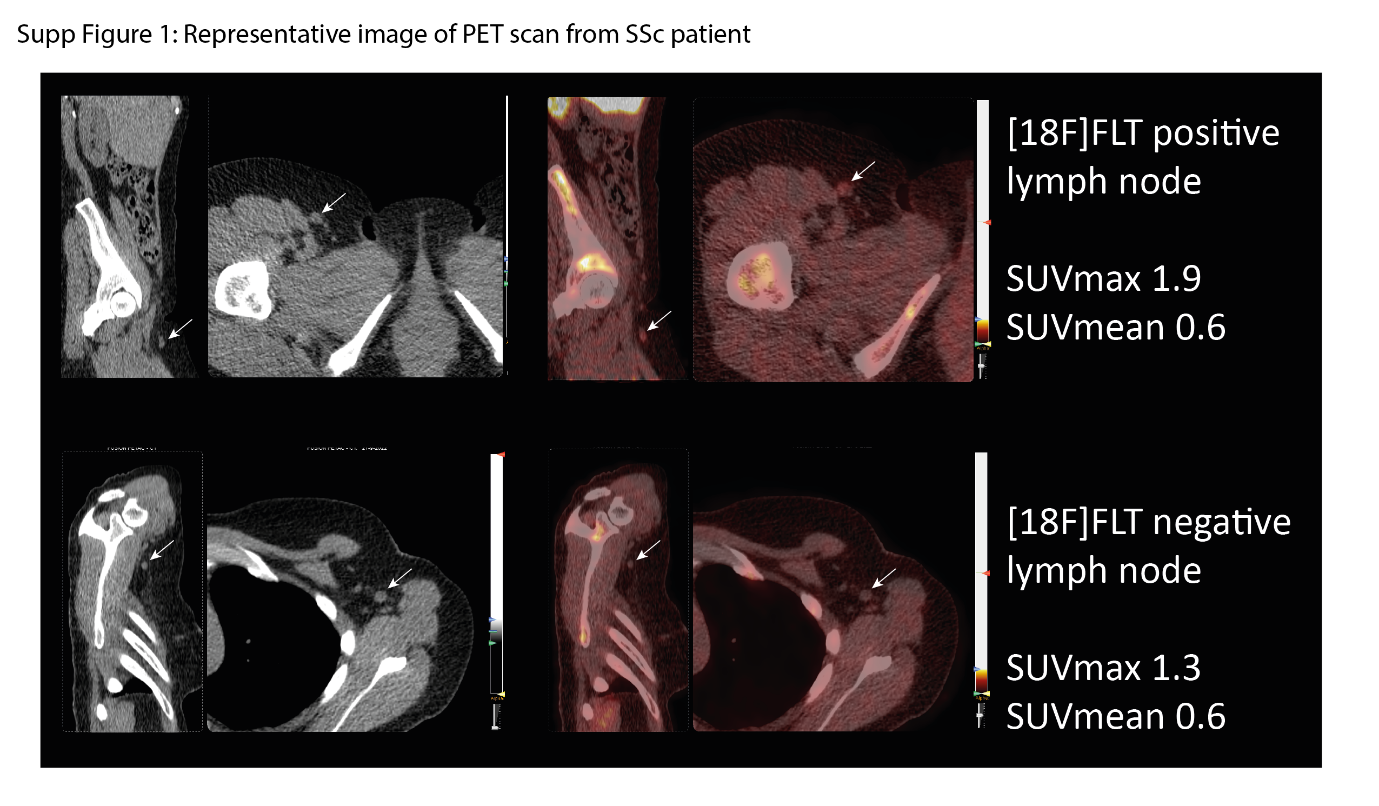

**Supplementary Figure 1. Representative image of [^18^F] FLT PET/CT scan** of a patient with SSc exhibiting with arrows detection, among the affected skin draining lymph nodes, of a FLT-positive (inguinal right) and a FLT-negative (axillar left) LN based on high or low [^18^F] FLT signal represented with SUV_mean_ and SUV_max_ values (g/ml).

**Supplementary Figure 2**

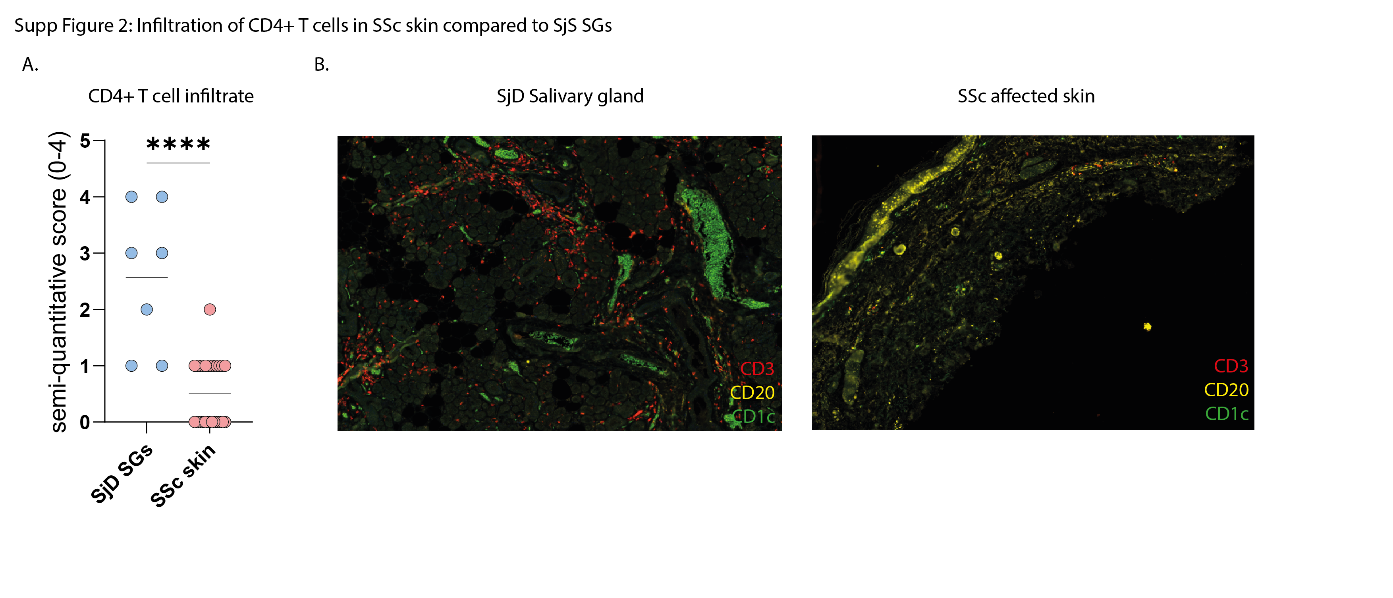
 **Supplementary Figure 2. Affected CTD of patients with SjD (salivary glands-SGs) exhibit higher infiltration of CD4+ T cell clusters compared to SSc affected skin.** (A) Semi-quantitative score of CD4+ T-cell infiltration within affected SjD salivary glands (SGs) (n=7) and SSc affected skin (n=20). (B) Representative multiplex immunofluorescence stainings exhibiting robust interaction between T-/B- cells withing APC rich areas in SjS SGs but not as prominent in SSc affected skin.

**Supplementary Figure 3**

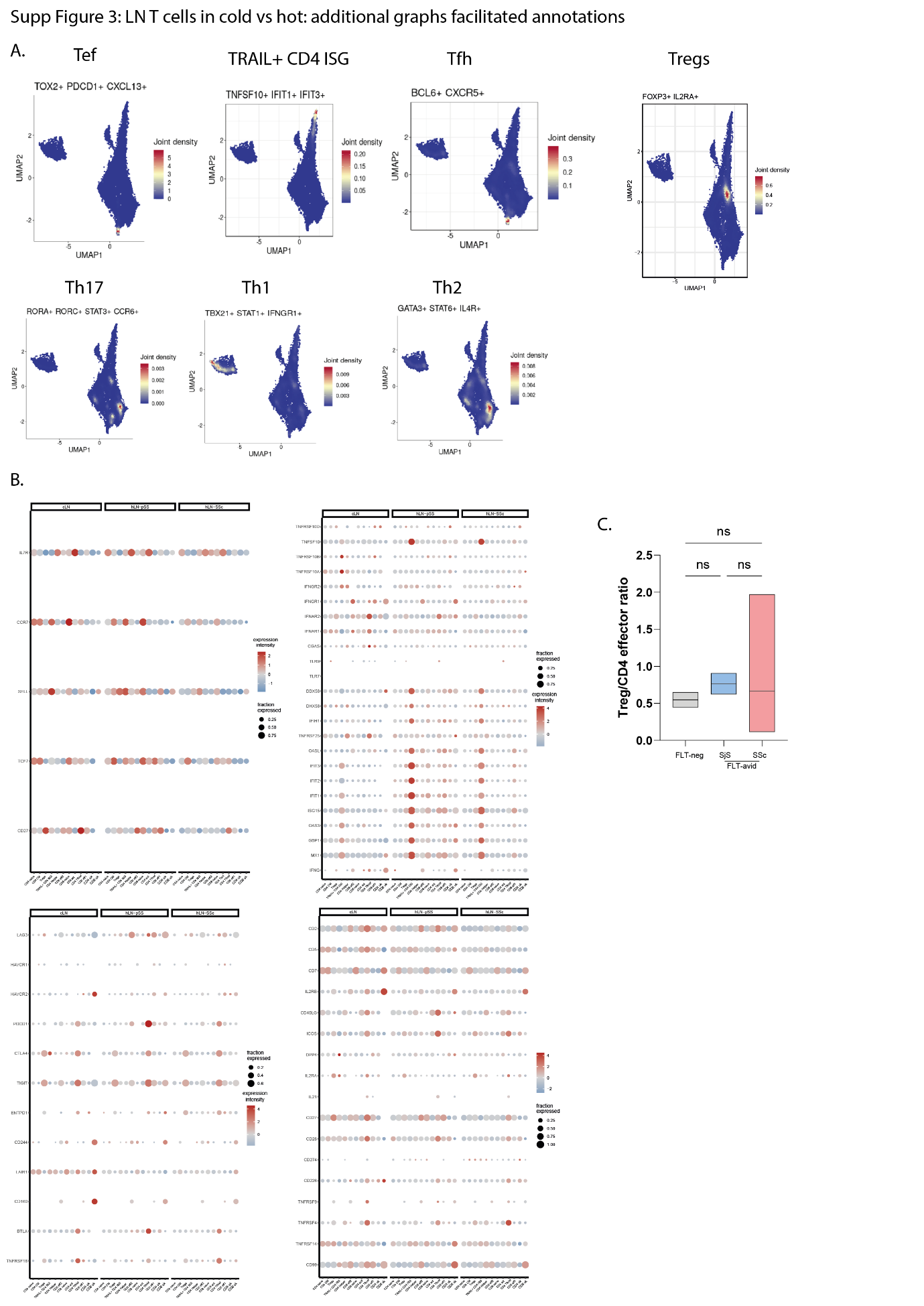

**Supplementary Figure 3. Additional single-cell RNA and CITE-seq analyses that facilitated annotation of CD4+ T-cell subsets in patients’ active and non-active lymph nodes.** (A) 2D Joint Density plots of selected markers reflective of Tph, TRAIL+ ISG, Tfh, Treg, Th17, Th1, Th2 specific gene signatures. (B) 2D dot plots showing the percentage and level of gene expression of selected type I IFN signaling, maturation, cytotoxic and activating/inhibitory receptor genes between T-cell clusters identified in active and non-active lymph nodes. (C) Cell ratio of Tregs/CD4 effector T cells between negative and positive LNs of patients with SjD and SSc.

**Supplementary Figure 4**

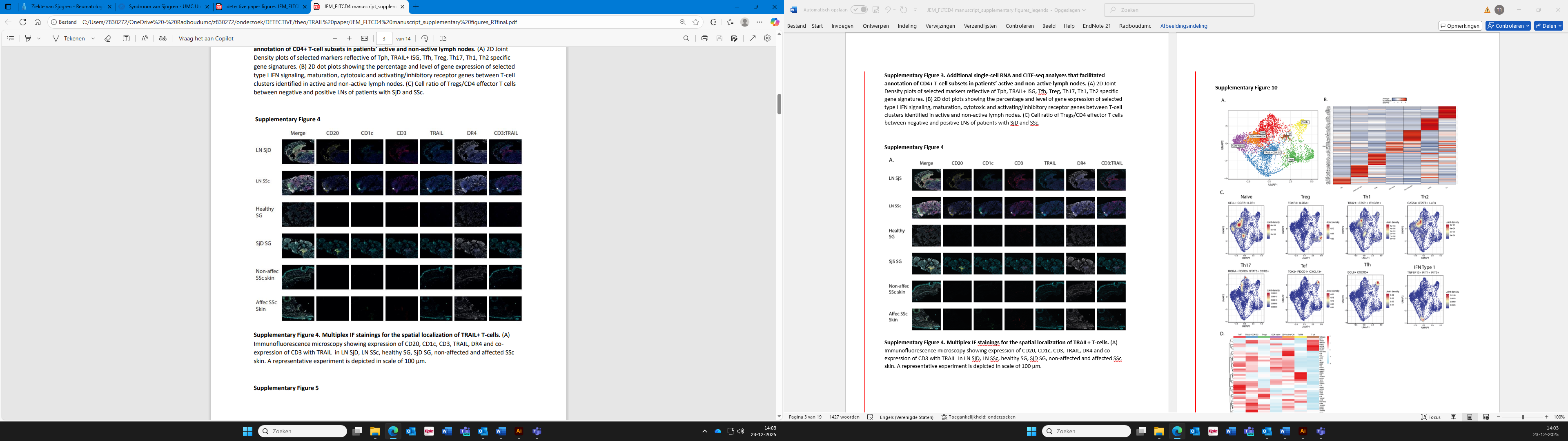

**Supplementary Figure 4. Multiplex IF stainings for the spatial localization of TRAIL+ T-cells.** (A) Immunofluorescence microscopy showing expression of CD20, CD1c, CD3, TRAIL, DR4 and co-expression of CD3 with TRAIL in LN SjD, LN SSc, healthy SG, SjD SG, non-affected and affected SSc skin. A representative experiment is depicted in scale of 100 µm.

**Supplementary Figure 5**

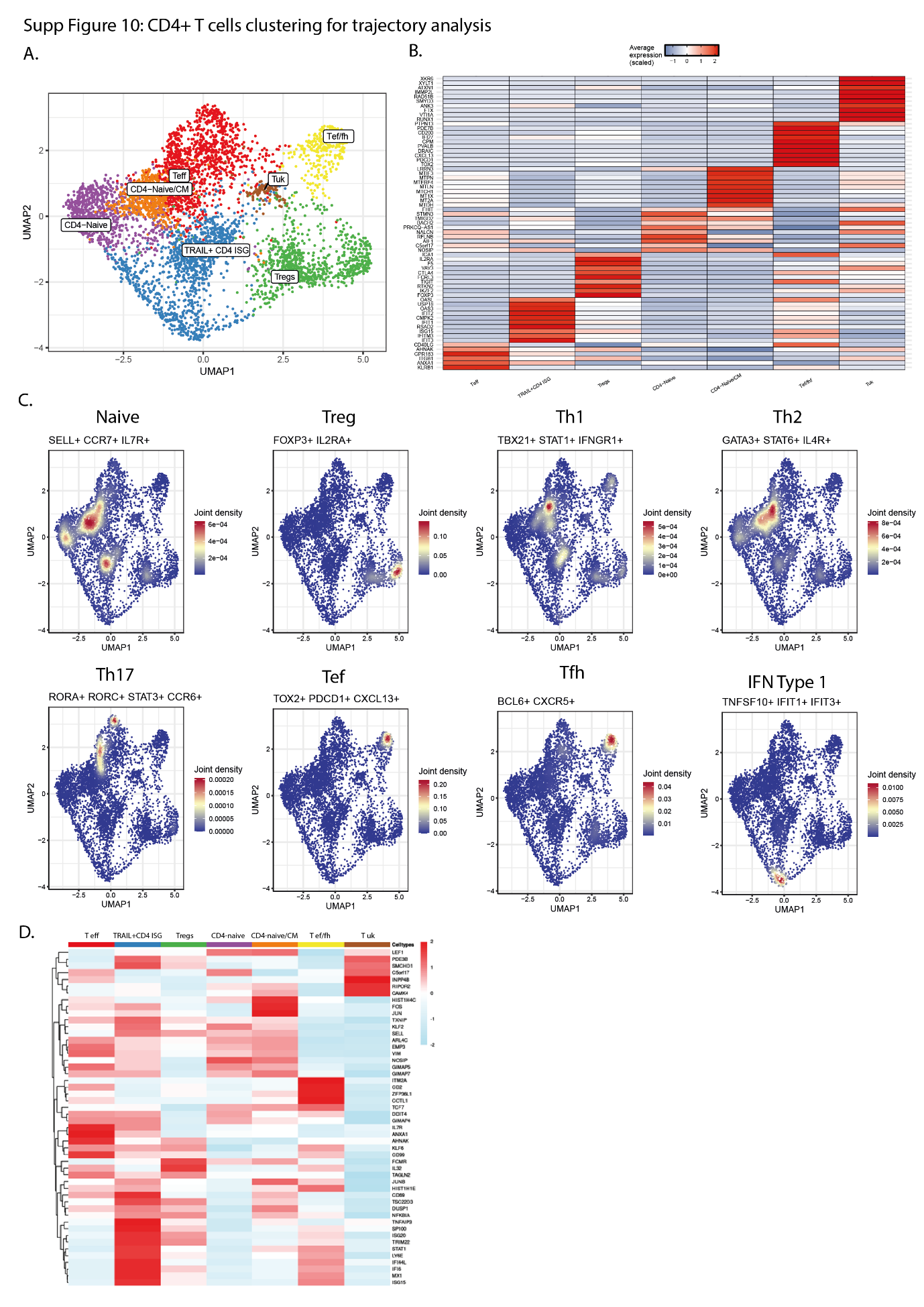

**Supplementary Figure 5. Focused subcluster single-cell RNA sequencing analysis of CD4+ T-cells to perform trajectory analysis.** (A) UMAP of the transcriptionally distinct CD4+ T-cell clusters in patients’ lymph nodes; CD4 naive, CD4 naive/CM, TRAIL+ CD4 ISG, T uk (unknown), Tregs, Tef/fh. (B) Heatmap illustrates the top 10 differentially expressed genes in each identified cell cluster of CD4+ LN T-cells. (C) 2D Joint Density plots of selected markers reflective of Naive, Treg, Th1, Th2, Th17, Tef, Tfh, IFN Type 1 specific gene signatures. (D) Differential dynamic genes (DYG) contributing to the clustering of each distinct CD4+ T-cell cluster.

**Supplementary Figure 6**

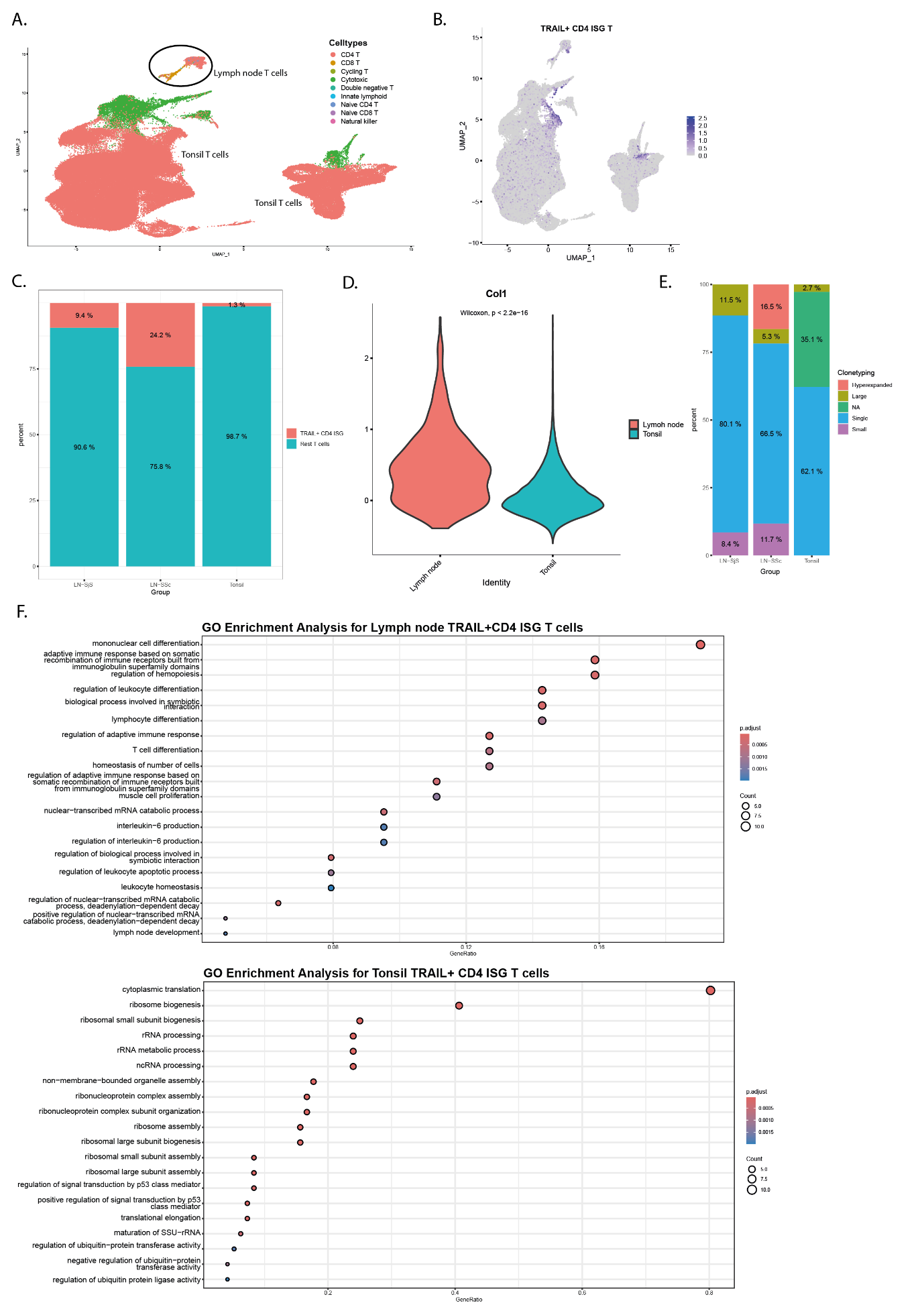

**Supplementary Figure 6. CD4+TRAIL+ ISG T cells exhibit unique expansion and immunoregulatory functional role within autoreactive lymph nodes.** For this analysis our single-cell RNAseq data from lymph node (LN) T cells of patients with SjS and SSc, were merged with single-cell RNAseq data of tonsils T-cells extracted from the atlas of human cells in the human tonsil [1] by Massoni-Badosa et al. (A) UMAP depicting the cell types present in the merged object of lymph node and tonsil T cells. (B) UMAP projection illustrating the presence and intensity of gene signature expression of TRAIL+ ISG CD4+ T cells. These cells were defined as expressing CD4, TNFSF10 and MX1 genes. (C) Bar graphs showing the frequency of TRAIL+ ISG CD4+ T cells between SSc/SjD reactive LNs and tonsils. (D) Violin plot comparing the intensity of ISG signature expression between LN and tonsil T-cells. (E) Bar graphs showing the percentage of expanded clones of TRAIL+ ISG CD4+ T cells between SSc/SjD reactive LNs and tonsils. (F-G) Gene ontology (GO) enrichment analysis based on the differentially expressed genes of TRAIL+ ISG CD4+ T cells in lymph node (top) versus tonsils (bottom).

**Supplementary Figure 7**

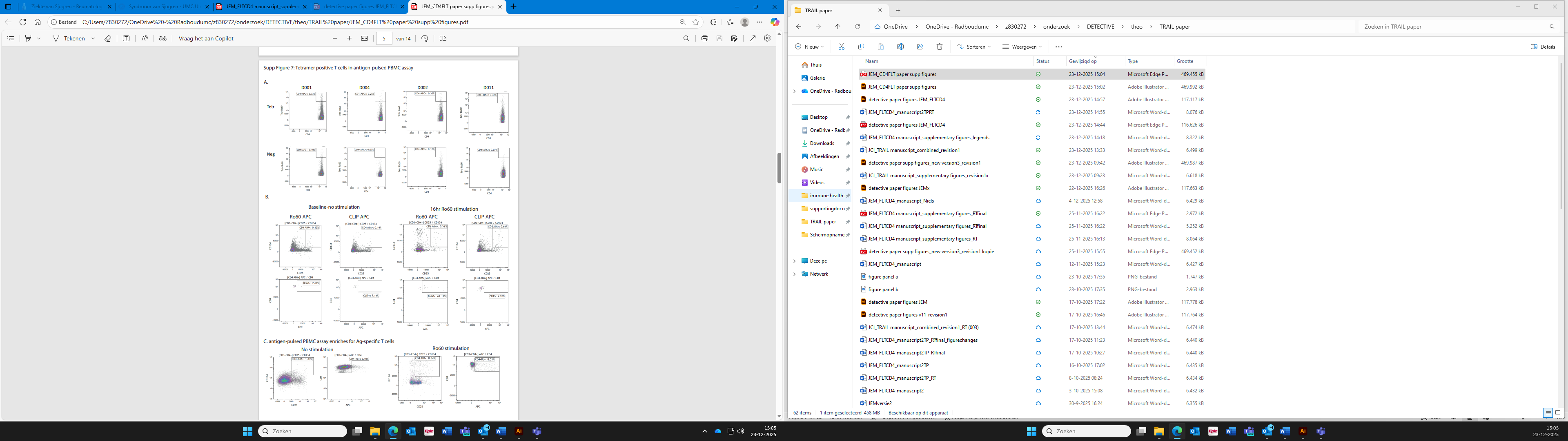

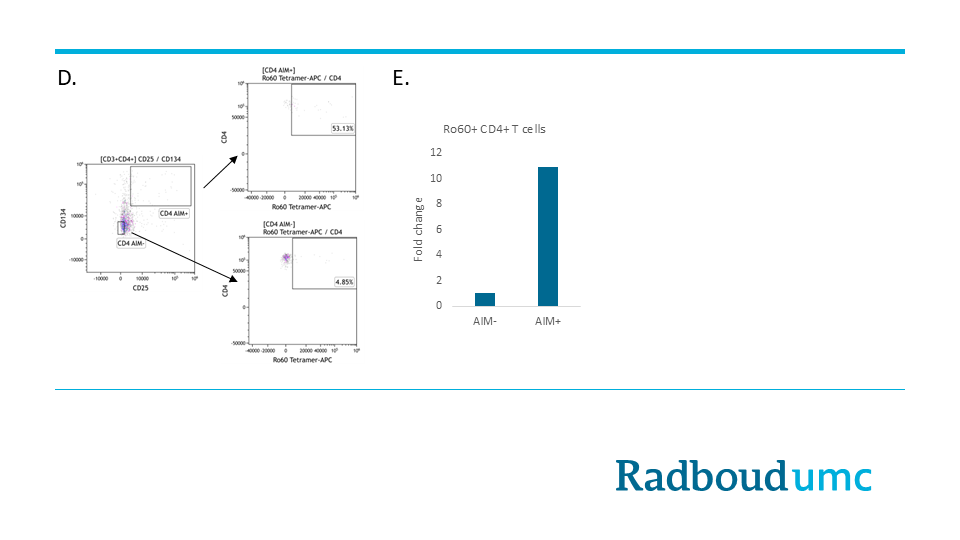

**Supplementary Figure 7. Detection of arCD4+ T-cells overlaps with detection of Ro60-specific T-cells with tetramers.** In 4 SjD patients with suitable HLA background, we performed an antigen-pulsed PBMC (Acitvation induced marker [AIM]) assay with the whole Ro60 antigen and combined with Ro60 (3 different peptides pooled) tetramer staining and found that the detection overlap between T-cell activation and tetramer staining was 60%. Furthermore, enrichment for antigen-responsive T-cells with the AIM assay also enhanced detection of autoreactive T-cells with tetramers and a similar percentage of T-cells were detected between the antigen-pulsed PBMC assay and tetramer staining (**Supp. figure 5C**). Importantly, within the AIM+ compartment there was an 11-fold increase in the presence of tetramer-specific T cells compared to the AIM- compartment (**Supp. figure 5D, E**). Collectively, these findings determine the validity of the AIM assay to detect autoreactive T-cells.

(A) Flow cytometry plots illustrate the detection of Ro60-specific CD4+ T-cells with the use of Ro60 tetramers conjugated with APC in 4 patients with SjD that are Ro60 seropositive. Results are compared with CLIP tetramers that were also conjugated with APC as negative controls. (B) Flow cytometric density plots of one representative experiment with PBMCs from SjD patient where tetramer staining in combination with the AIM assay was performed. An enriched number of Ro60-specific T-cells is detected at similar levels (C) with T cell activation markers and tetramer staining after 16-hour incubation of PBMCs with recombinant Ro60 protein. The overlap between activated T cells and tetramer+ cells is about 60%. (D) FACs plots showing the number of tetramer+ CD4+ T cells that are based on activation markers in the AIM- versus AIM+ cell compartment. (E) Fold change showing increased percentage of tetramer+ T cells in AIM+ compared to AIM- T cells.

**Supplementary Figure 8**

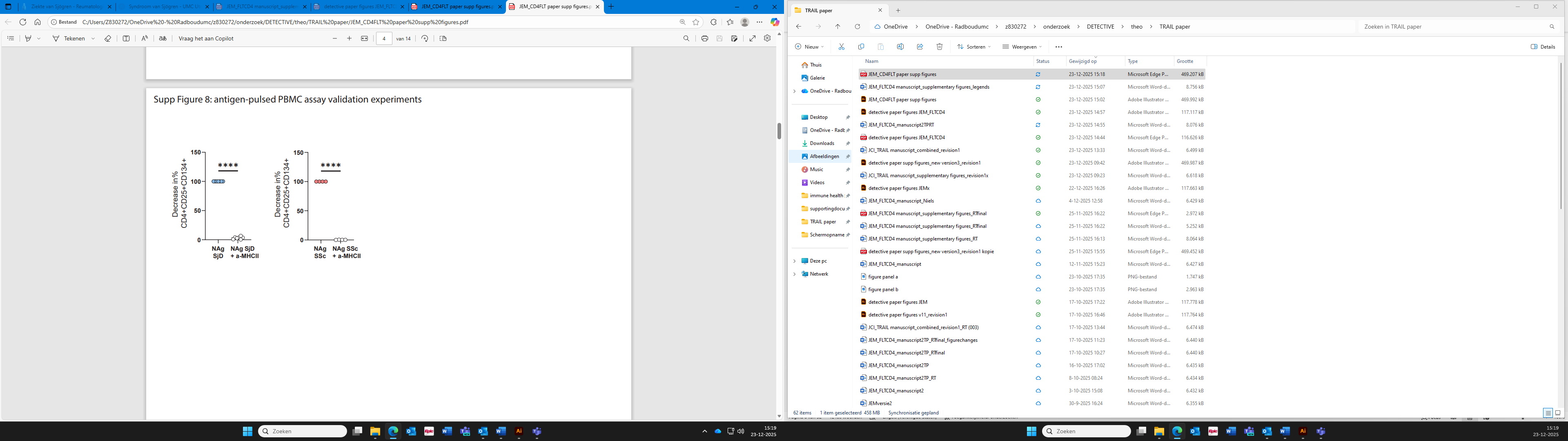

**Supplementary Figure 8. arCD4+ circulating T-cells are expanded in SjD and SSc patients in an MHCII-dependent manner.** Normalized decrease in % of CD4+AIM+ (CD4+CD25+CD134+) response under Scl70 or Ro60/La stimulation and blockage or not of MHC II interactions with anti-HLA-DR/DQ/DP moAb.

Statistics: in (A) RM one-way ANOVA, with Dunnett’s multiple comparisons test, *p<0.05, **p<0.01; in (C) two-tailed paired t-test, ****p<0.0001; in (E) ordinary one-way ANOVA, with Tukey’s multiple comparisons test, *p<0.05, ***p<0.001;

**Supplementary Figure 9**

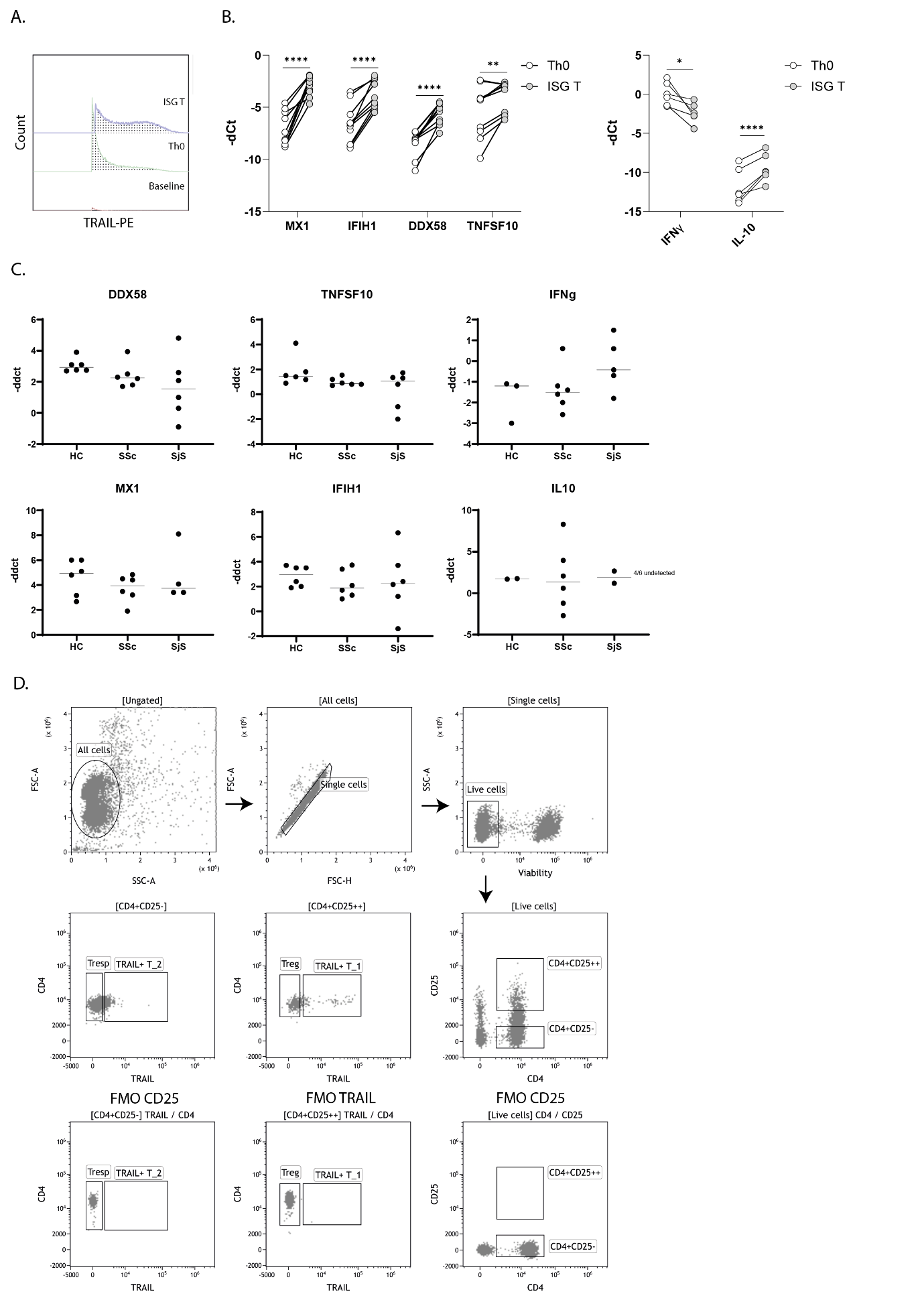

**Supplementary Figure 9.** (**A)** Overlay histograms of one representative experiment illustrating expression of TRAIL in baseline, Th0 and ISG differentiated T-cells. **(B)** Relative gene expression (−dCt) of the depicted genes measured with qPCR on Th0 or ISG differentiated T-cells (n=4). (**C**) Relative gene expression (-ddct) of the depicted genes comparing ISG T/Th0 differentiated cells from healthy controls (HC) and patients with SSc and SjS (n=6 per group). (**D**) Gating strategy that was used to sort TRAIL+ T cells (CD4+CD25-TRAIL-), Tregs (CD4+CD25++TRAIL-) and Tresp(onsive) cells (CD4+CD25-TRAIL-) for the CD4 effector proliferation suppression assay.

Statistics for panel B were performed with ordinary one-way ANOVA with Tukey’s multiple comparisons test, *p<0.05.

**Supplementary Figure 10**

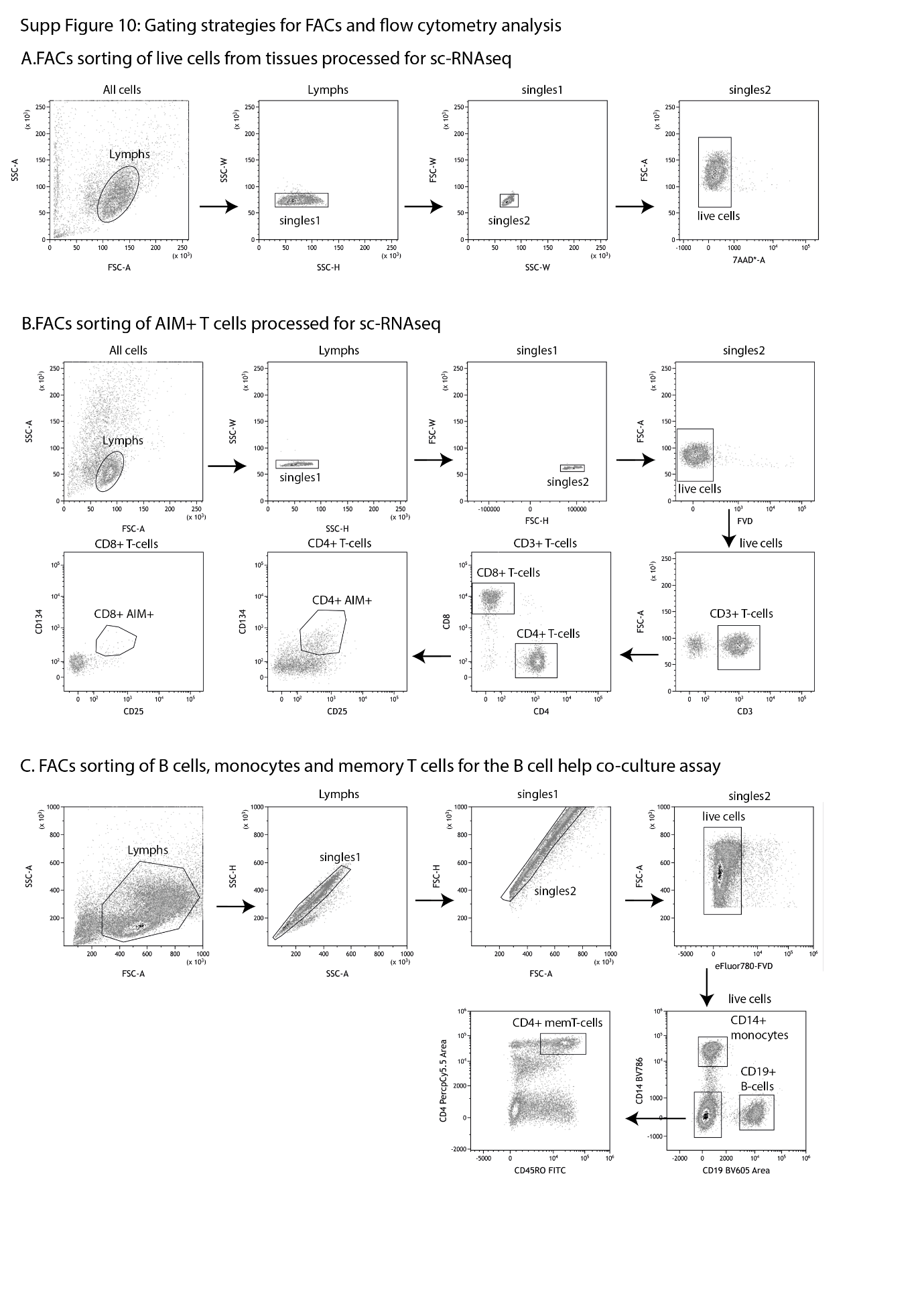

**Supp Figure 10. Gating strategies exhibiting fluorescently activated cell sorting (FACs)** of (**A**) Viable lymph node lymphocytes that were subsequently processed for 10x single-cell RNA sequencing. (**B**) Viable AIM+ CD4+ T-cells and CD8+ T-cells that were subsequently processed for 10x single-cell RNA sequencing. (**C**) B-cells, monocytes and memory T-cells that were subsequently co-cultured in vitro for the B-cell help immunoassay.

**Supplementary Figure 11**

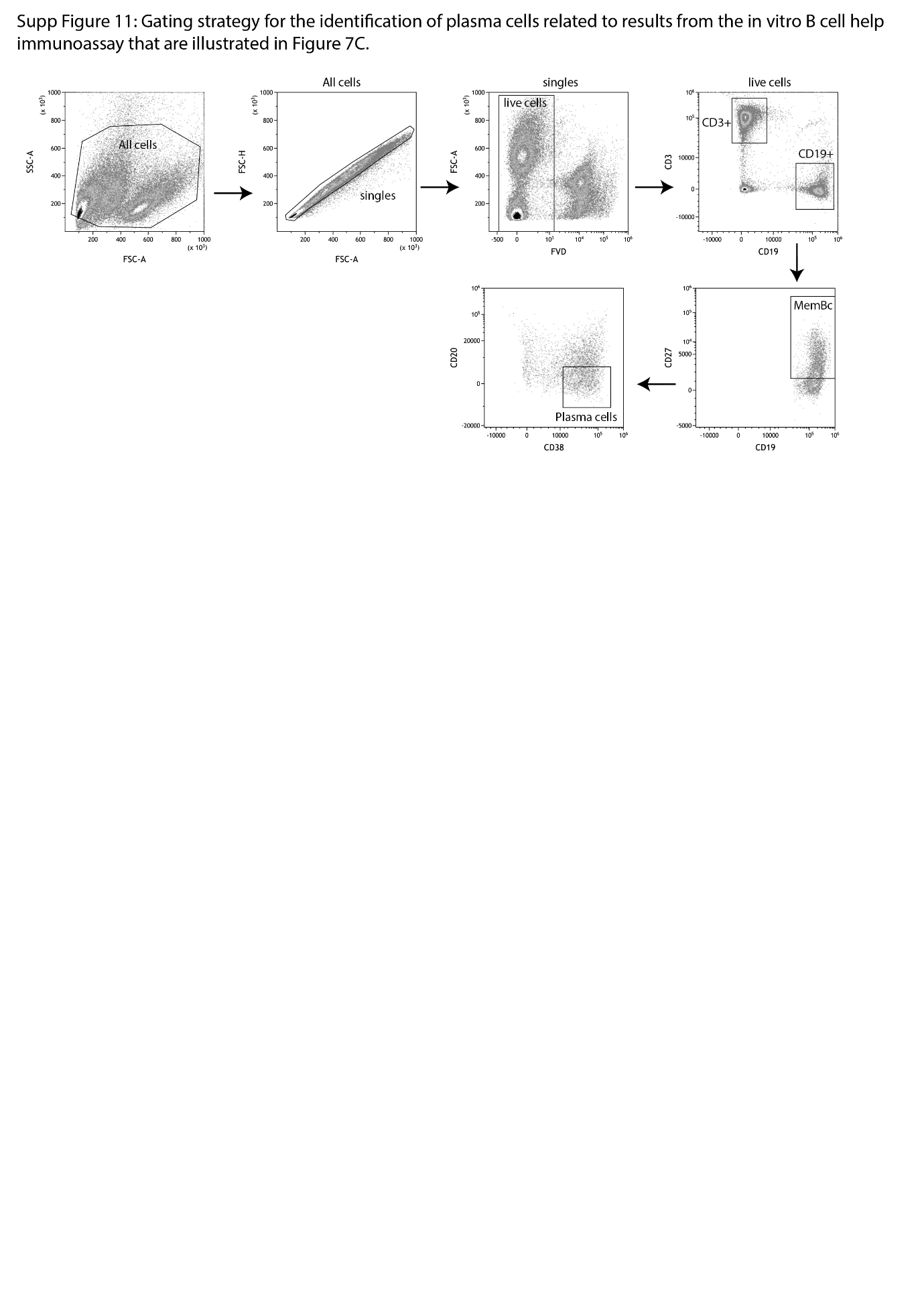

**Supplementary Figure 11. Gating strategy for the identification of plasma cells related to results from the in vitro B-cell help assay that are illustrated in Figure 8.**
